# Spatio-temporal, optogenetic control of gene expression in organoids

**DOI:** 10.1101/2021.09.26.461850

**Authors:** Ivano Legnini, Lisa Emmenegger, Alessandra Zappulo, Ricardo Wurmus, Anna Oliveras Martinez, Cledi Cerda Jara, Anastasiya Boltengagen, Talé Hessler, Guido Mastrobuoni, Agnieszka Rybak-Wolf, Stefan Kempa, Robert Zinzen, Andrew Woehler, Nikolaus Rajewsky

**Affiliations:** Laboratory for Systems Biology of Gene Regulatory Elements, Berlin Institute for Medical Systems Biology (BIMSB), Max Delbrück Center for Molecular Medicine (MDC) in the Helmholtz Association, Hannoversche Str. 28, 10115 Berlin; Systems Biology of Neural Tissue Differentiation, Berlin Institute for Medical Systems Biology (BIMSB), Max Delbrück Center for Molecular Medicine (MDC) in the Helmholtz Association, Hannoversche Str. 28, 10115 Berlin; Bioinformatics and Omics Data Science, Berlin Institute for Medical Systems Biology (BIMSB), Max Delbrück Center for Molecular Medicine (MDC) in the Helmholtz Association, Hannoversche Str. 28, 10115 Berlin; Systems Biology Imaging, Berlin Institute for Medical Systems Biology (BIMSB), Max Delbrück Center for Molecular Medicine (MDC) in the Helmholtz Association, Hannoversche Str. 28, 10115 Berlin; Proteomic and Metabolomics Platform, Berlin Institute for Medical Systems Biology (BIMSB), Max Delbrück Center for Molecular Medicine (MDC) in the Helmholtz Association, Hannoversche Str. 28, 10115 Berlin; Organoid platform, Berlin Institute for Medical Systems Biology (BIMSB), Max Delbrück Center for Molecular Medicine (MDC) in the Helmholtz Association, Hannoversche Str. 28, 10115 Berlin

**Keywords:** Organoids, optogenetics, spatial transcriptomics, single-cell transcriptomics, CRISPR/Cas, neurodevelopment, Sonic Hedgehog (*SHH*)

## Abstract

Organoids derived from stem cells become increasingly important to study human development and to model disease. However, methods are needed to control and study spatio-temporal patterns of gene expression in organoids. To this aim, we combined optogenetics and gene perturbation technologies to activate or knock-down RNA of target genes, at single-cell resolution and in programmable spatio-temporal patterns. To illustrate the usefulness of our approach, we locally activated Sonic Hedgehog (*SHH*) signaling in an organoid model for human neurodevelopment. High-resolution spatial transcriptomic and single-cell analyses showed that this local induction was sufficient to generate stereotypically patterned organoids in three dimensions and revealed new insights into *SHH*’s contribution to gene regulation in neurodevelopment.

With this study, we propose optogenetic perturbations in combination with spatial transcriptomics as a powerful technology to reprogram and study cell fates and tissue patterning in organoids.

## Introduction

Organoid culture has proven to be a transformative technology by offering the opportunity to access unique features of human development and to model complex attributes of human disease *in vitro*. Organoid development relies on the intrinsic property of stem cells to differentiate and self-organize in three-dimensional space (Lancaster et al., 2013). Specific developmental trajectories can be further promoted by treatment with molecules controlling cell differentiation or by genetic manipulations, and more complex architectures can be achieved by fusing organoids with different properties into assembloids (Kelley and Pasca, 2022). Controlling organoid tissue patterning in a programmable manner and with spatio-temporal resolution, however, remains an unmet challenge.

On the other hand, methods such as single-cell RNA sequencing and spatial transcriptomics have proven immensely useful to describe the molecular signatures of cell states and their relationships within a tissue, both in health and disease. However, to interrogate and elucidate the molecular mechanisms that explain these data, it is essential to perturb gene expression, ideally at single-cell resolution and conditionally in both space and time – targeting a live cell or even different live cells of interest within the tissue (for example in different spatial positions) simultaneously and at a controlled time point.

To address both the need to control organoid patterning as well as the need for temporally controlled and spatially multiplexed (“programmable”) gene expression perturbations, we developed a flexible system that allows light-inducible activation and repression of target genes, by combining optogenetic transcription (Nihongaki et al., 2017, Yamada et al., 2018, De Santis et al., 2021) with CRISPR/Cas13 knock-downs (Abudayyeh et al., 2017, Cox et al., 2017, Konerman et al., 2018). We then engineered laser scanning and digital light projection setups for live-cell photo-stimulation and imaging and were able to “print” complex patterns of gene expression onto single cells and organoids.

To show that our approach can indeed control organoid patterning, we chose to locally activate the Sonic Hedgehog (*SHH*) pathway in an organoid model of human neural tube development, and quantified the consequences of this approach at single-cell resolution and with high-resolution spatial transcriptomics. *SHH* is a well-studied morphogen that is central to a variety of biological processes, including the dorsal-ventral patterning of the vertebrate neural tube (Ribes and Briscoe, 2009). We developed neural organoids to mimic the neural tube patterning and we optogenetically induced *SHH* in a pole of these organoids. Spatial transcriptomics, including both a standard low-resolution but unbiased “Visium” approach as well as multiplexed in-situ hybridization of 88 genes, including a custom-designed panel of genes linked to *SHH* biology at single-molecule resolution, revealed that localized *SHH* activation was sufficient to establish robust and well-defined gene expression territories, resembling those found in the ventral regions of the neural tube *in vivo*. Single-cell analysis allowed to more comprehensively reconstruct dorsoventral identities corresponding to the neural tube patterning and revealed novel insights into the impact of *SHH* signaling on the differentiation of neuronal progenitors, for example the strong activation of IGF pathway modulators, the differentiation of cells expressing pericyte markers and the spatial modulation of axon guidance molecules.

## Results

### Light-inducible gene activation and knock-down

In order to perturb RNA expression with spatial resolution, we adopted, constructed and optimized a variety of tools based on the combination of optogenetic protein elements to allow spatial control by photo-stimulation, with gene perturbation effectors to induce gene activations and/or knock-downs. The general design that we adopted consists of an activation module, based on light-inducible CRISPR/Cas9, Tet-ON or Cre/Lox systems, which can be used to switch on endogenous promoters or exogenous expression cassettes, and a CRISPR/Cas13 module, which can be coupled with the activation module for knocking down transcripts of interest.

To optogenetically induce gene activations, we first used the <u>s</u>plit <u>C</u>RISPR-Cas9-based <u>P</u>hotoactivatable <u>T</u>ranscription <u>S</u>ystem (*SCPTS*, Nihongaki et al., 2017), which consists of an enzymatically dead Cas9 (dCas9) split into N- and C-terminal domains and fused to the photoinducible dimerization moieties pMag and nMag. Blue light triggers pMag-nMag dimerization, thereby reconstituting dCas9, that incorporates available guide RNAs to target an intended promoter. The SCPTS system activates transcription nearby (CRISPRa), via fused or associating activation domains (VP64, p65 and HSF1, Fig. 1a). This system has been shown to be a potent transcriptional activator under blue light illumination (Nihongaki et al., 2017). We established the SCPTS system in transfected HEK cells using a custom programmable LED board (Methods), which accommodates a 96-well cell culture plate and that can be used in a cell culture incubator. Transcriptional induction levels were comparable to Nihongaki et al. (2017, Fig. S1a).

**Figure 1.**
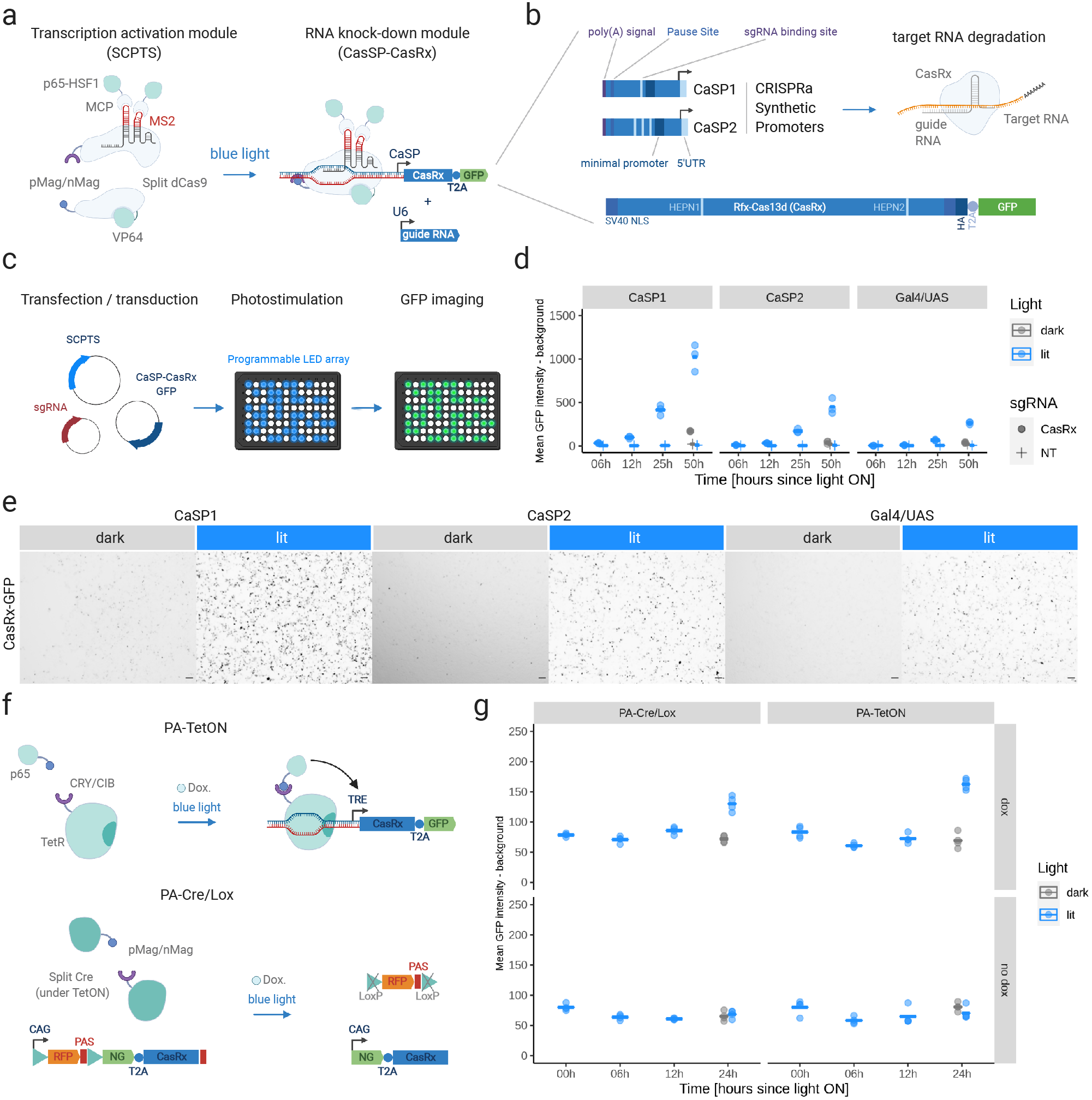
Light-inducible gene activation and knock-down modules. **a.** Left: light-inducible transcription activation module (SCPTS, Nihongaki et al., 2017). The Cas9 sgRNA is shown associated to the N-ter part of dCas9 in the dark state. Right: RNA knock-down module: CasRx transcription is driven by a synthetic promoter controlled by the SCPTS module. A guide RNA for targeted knock-downs can be co-expressed from a U6 promoter. **b.** Synthetic promoters for light-inducible transcription of CasRx (CaSP1/2), containing upstream elements for reducing spurious transcription (poly(A) site, pause site), a minimal CMV promoter containing TFIIB binding site/TATA box, an initiator and synthetic 5’UTR, one or three sgRNA binding sites. The CasRx cassette (below) contains a T2A-GFP tag, two nuclear localization signals (NLS) and an HA tag, as in Konermann et al. (2018). HEPN1-2 (Higher Eukaryotes and Prokaryotes Nucleotide-binding) catalytic domains are indicated. **c.** Experimental setup: HEK293T cells are transfected in a 96-well plate, which is placed on a LED-board for blue-light stimulation. At each time point, cells are imaged for GFP quantification. **d.** For the SCPTS system, background-subtracted mean GFP intensity is plotted for the selected time points (dark or lit), with one of the three promoters (CaSP1/2, Gal4/UAS), with either a non-targeting guide (NT) or the CasRx promoter-targeting guide (CasRx). Horizontal bars: mean of all replicates (three). **e.** Representative images for the SCPTS CasRx system with the three promoters in presence of the CasRx sgRNA. Scale bar: 100 μm. Images were taken at 24h post-transfection with a Keyence BZ-X710 with 4x magnification, in independent experiments from those quantified in panel d. **f.** Schematic representation of: a PA-TetON system consisting of a TetR/p65 transactivator fused with CRY/CIB photodimers; a PA-Cre/Lox based on a split Cre fused with pMag/nMag photodimers. The PA-TetON system is coupled with a Tet-Responsive Element (TRE) promoter controlling CasRx expression, the PA-Cre/Lox system is combined with a LoxP-RFP-LoxP-CasRx cassette, which places CasRx under a CAG promoter upon the Cre-mediated removal of the RFP cassette. **g.** Same as d., for the PA-TetOn and PA-Cre/Lox systems, over a 0-24h timecourse, with or without doxycycline treatment (dox/no dox).

We then tested a promoter/sgRNA pair, previously used in a similar context (Gal4/UAS, Nihongaki et al, 2015a), and additionally designed two synthetic promoters (CRISPRa Synthetic Promoter – CaSP1 and 2, partially based on Loew et al., 2010, Fig. 1b) to drive transcription of any given expression cassette. In this case, we activated the expression of a GFP-tagged CasRx, which can be further used for targeted RNA knock-downs. We transfected HEK cells with the plasmids encoding the transcription system, the sgRNA and a CasRx-T2A-GFP cassette under the control of one of the three promoters and imaged GFP over time upon photo-stimulation (Fig. 1c, S1b). Both synthetic promoters CaSP1 and CaSP2 were more active than the UAS promoter (Fig. 1d). CaSP1 induced a ~45-fold-change activation after 50h illumination over the non-targeting guide control, with ~16% leakage in the dark. In contrast, CaSP2 elicited a ~21-fold induction, with a leakage of ~9%. A constitutive CasRx-T2A-GFP cassette under the control of a strong EF1a promoter produced GFP with no substantial difference between light and dark conditions (Fig. S1c). Representative images of the SCPTS/Cas13 photo-stimulation are shown in Fig. 1e.

In search for alternative means of light-inducible gene activation with different properties, we also used a light-inducible Tet-ON transcription system (Yamada et al., 2018) and a light-inducible Cre/Lox system (De Santis et al., 2019), which both require a double light and doxycycline switch (Fig. 1f). For the Tet-ON system (Yamada et al., 2018), we cloned a CasRx-T2A-GFP cassette under a Tet Responsive Element (TRE) promoter and transduced HEK293T cells with two lentiviruses expressing the components of the transcription system and the inducible CasRx. For the Cre/Lox system, we used a double Piggybac system to generate a stable line, with one transposon carrying the split Cre (under a Tet-ON promoter) and the other carrying a LoxP-RFP-LoxP cassette controlled by a CAG promoter, followed by NeonGreen-T2A-CasRx, which is repositioned under the promoter upon Cre activation. Both these systems proved reasonably tight and light-responsive, with circa two-fold activation over the background and no-doxycycline control at 24h after the light stimulus (Fig. 1g). PA-TetON was therefore considerably weaker than the SCPTS CaSP1-2 systems, which on the other hand were transiently transfected and therefore present in high copy number. As the Cre/Lox system is tagged with a different fluorescent protein (NeonGreen vs EGFP), it was not possible to directly compare it with the other systems. Microscopy-based quantifications were additionally confirmed by flow cytometry measurements (Fig. S1d-g).

An extensive report of the knock-down module efficacy and additional attempts at designing light-inducible Cas13 proteins are reported in a Supplementary Note and in Figures S2-3.

### Spatial programming of optogenetic stimulations

In order to leverage the potential of optogenetic RNA perturbations, we not only need programmable gene activation and knock-down modules, but also the means to program them spatially. To this end, we tested three different approaches (Fig. 2a), including photomasks, directed laser stimulations and a programmable Digital Micromirror Device (DMD).

**Figure 2.**
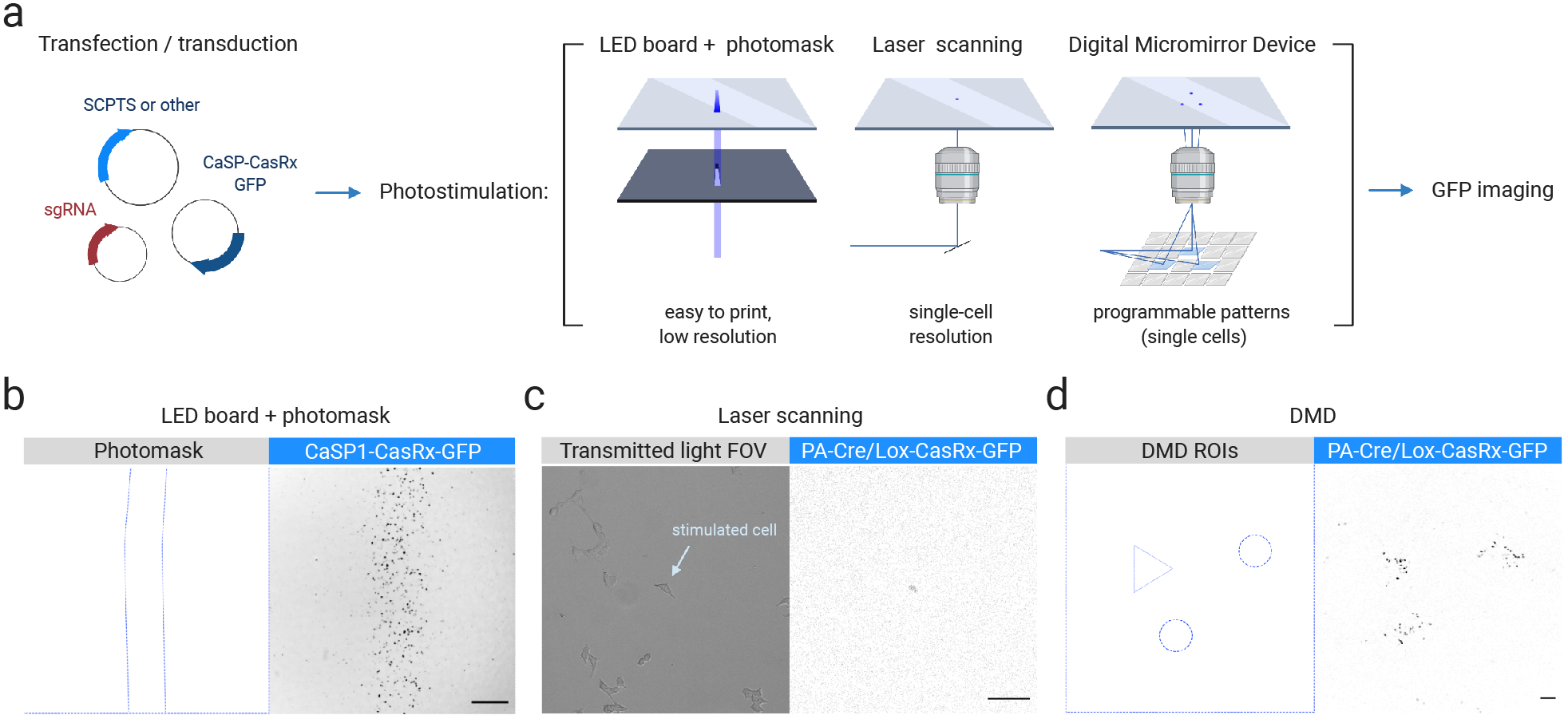
Spatial programming of optogenetic stimulations. **a.** Spatially induced photo-stimulations: cells are stimulated with a blue LED array combined with a black photomask, or with a confocal microscope setup by laser scanning, or with a Digital Micromirror Device microscope by a LED source projected through a micromirror array to the cell culture plate bottom. **b.** Representative fluorescent microscope image of a photomask stimulation of the CaSP1-CasRx system in HEK cells. Blue: photomask shape. Signal is GFP in grey scale. Scale bar: 500 μm. **c.** Representative image of a single-cell Cre/Lox CasRx stimulation in HEK cells, performed with 100 Hz laser scanning within a confocal microscope setup. Left: transmitted light image, right: confocal image of the GFP signal in grey scale. Scale bar: 100 μm. **d**. Representative image of a complex pattern stimulation of the Cre/Lox CasRx system in hiPSCs performed with the DMD setup. The selected ROIs are shown on the left, while the NeonGreen signal imaged with a confocal setup after 24h stimulation within the DMD setup is shown in grey scale on the right. Scale bar: 100 μm.

First, we applied a printed photomask between cells that express light-inducible CasRx cassettes and the light source (Fig. 2b). This setup is simple to construct, as a laser printer is sufficient to generate a negative stencil which is then attached beneath a cell culture plate. The plate is then placed on top of a LED array and light stimulation is provided for the desired time. The array is capable of generating complex patterns of activation with a resolution on the order of hundreds of micrometers, provided by the photomask (Fig. 2b). However, we noticed some diffused induction beyond the edges of the photomasks, possibly due to reflection and refraction within the plate material. This approach, while simple and intuitive, provides a narrow application spectrum and is largely limited to thin 2D cultures.

We next tested if we could achieve precise spatial activation of gene expression at single-cell resolution. We employed a laser scanning-based photo-stimulation approach, and we performed live-cell imaging with a laser scanning confocal microscope. We induced CasRx expression from the light inducible Cre/Lox system by scanning a region of interest (ROI) containing a single-cell. Using this approach, we successfully stimulated a single cell in a field of view containing several cells. (Fig. 2c). This approach, though slightly more difficult to implement, can be applied to more complex structures and tissues as the laser can be focused on specific cells of interest.

Lastly, we constructed a DMD microscope, combined with a cell culture chamber for live-cell stimulation and imaging. The DMD is controlled by a simple micromanager/ImageJ-based graphical user interface (GUI) which allows to intuitively program spatial activation patterns. We describe the details for building and programming this “point-and-shoot” setup in the methods section. Within the GUI, we can draw multiple ROIs with varying complexity directly onto the sample, which will then be illuminated by the DMD for the desired time intervals and intensities (Fig. 2d).

### Optogenetic stimulation of the SHH pathway in hiPSCs

To test the ability of our setup to perturb biologically relevant processes, we focused on the induction of Sonic Hedgehog signaling in stem cells first (Fig. 3a), and to pattern neural organoids later. We first designed three sgRNAs for activating the *SHH* promoter with the SCPTS system (Fig. S4a). We transfected HEK293T cells with all the SCPTS modules and each sgRNA, then quantified *SHH* mRNA expression after 24 hours of photo-stimulation. Guide 1 was the most efficient, increasing *SHH* mRNA expression ca. 800-fold over a non-targeting guide (Fig. S4b). In addition, we designed guides for another morphogen involved in neurodevelopment, *BMP4* (Fig. S4c). Guide 3 was the most effective, inducing a 4-fold increase in *BMP4* mRNA levels (Fig. S4d). Leakage was approximately 5% for *SHH* guide 1 and 30% for *BMP4* guide 3. We note that *BMP4* is natively expressed in HEK cells, which is likely a reason for the lower induction and higher leakage. We assessed whether the induced *SHH* exerted its biological activity by stimulating the expression of its targets, upon transfection of the SCPTS system in hiPSCs. We measured *FOXA2*, *FOXG1*, *NKX2-1*, *NKX6-2* and *OLIG2* expression after 24, 48 and 72 hours of stimulation. *SHH* reached its highest level of activation at 24 hours and decreased at 48 and 72 hours (Fig. S4e-f). Within 72 hours of light stimulation, *FOXA2, NKX6-2* and *OLIG2* were significantly upregulated in neural induction media, but not in stem cell media (Fig. S4e-f). Light-inducible activation of the *SHH* promoter is therefore sufficient to stimulate *SHH* transcription and exert a detectable biological effect by inducing the expression of some of its known targets. In parallel, we used the PA-Cre/Lox system to generate a stable hiPSC line that overexpresses a NeonGreen-SHH cassette upon light stimulation and doxycycline treatment (SHH-GFP for simplicity, De Santis et al., 2021, suppl. video 1). With this system, we observed stronger *SHH* mRNA induction upon light stimulation but also higher leakage, and the same was true for its targets (Fig. S4e-g). Robust expression of *FOXA2* was visible at the protein level after inducing *SHH* for 6-7 days in defined ROIs with the DMD setup (Fig. 3b).

**Figure 3.**
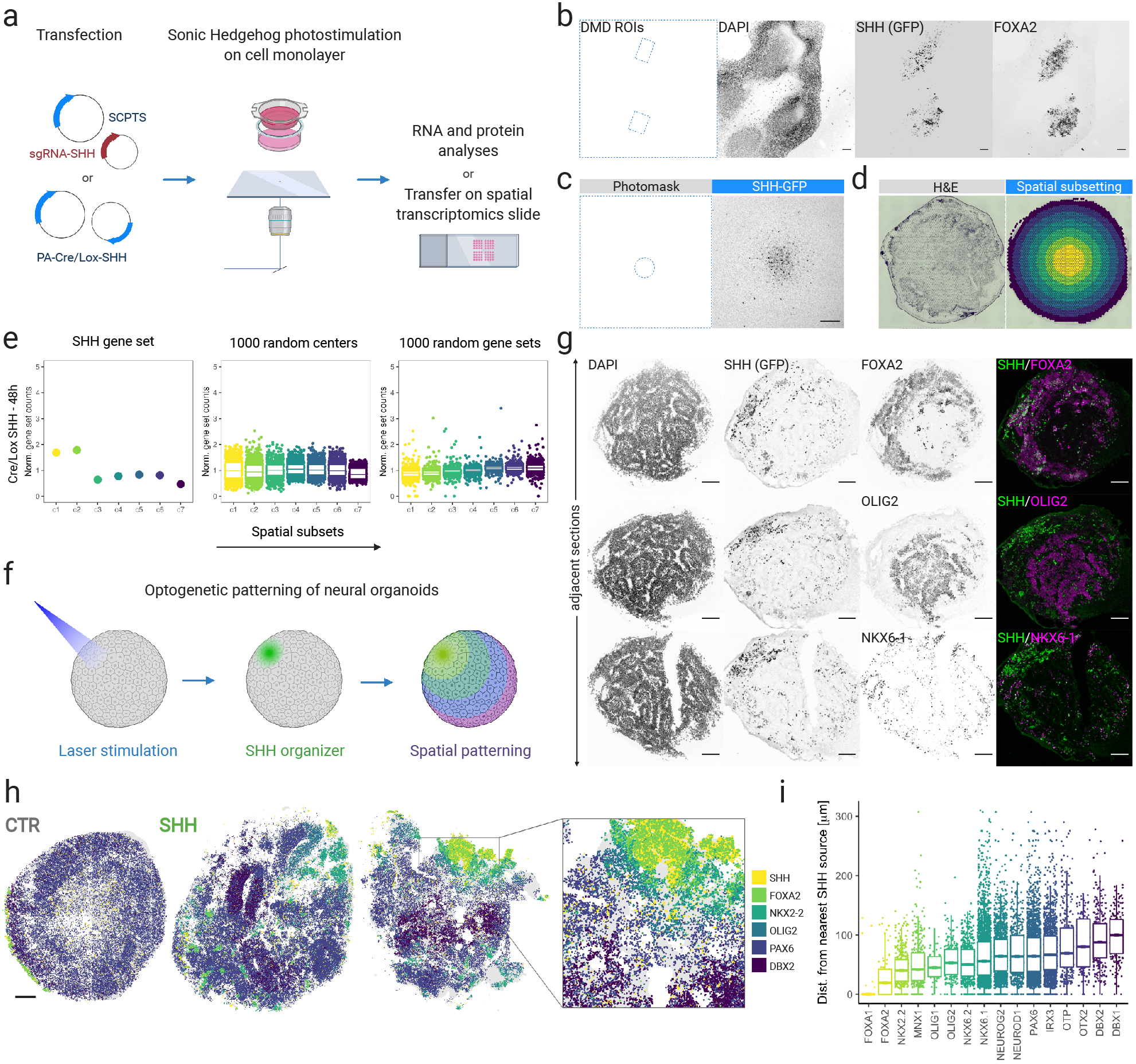
Optogenetic stimulation of Sonic Hedgehog in human stem cells and organoids. **a.** Coupled optogenetic stimulation of *SHH* and spatial readouts. **b.** Imaging of DAPI, SHH-expressing cells marked by NeonGreen (labeled as GFP for simplicity) and *FOXA2* (immunofluorescence) in hiPSCs 6 days post-stimulation. *SHH* was induced in two ROIs (left) with the DMD setup. Signal is shown in greyscale for each channel. Scale bar: 100 μm. **c.** Representative image of hiPSCs Cre/Lox *SHH* cultured as a monolayer on a PET membrane, induced in the center with a 500 μm-wide circular photomask (left). Signal is shown in greyscale (right). Scale bar: 500 μm. **d.** Left: representative H&E staining of a hiPSC layer cultured on a membrane and transferred onto a VISIUM slide. Right: representative spatial subsetting of spots within a capture area of a VISIUM slide into 7 concentric circles, centered on the SHH-induced area. **e.** Left: normalized counts of a *SHH* gene set (*SHH*, *FOXA2*, *FOXG1*, *NKX2-1*, *NKX2-2*, *NKX6-2*, *NKX6*-1, *OLIG2*) in the 7 concentric circles c1-7, color coded as in d, in hiPSCs stimulated for 48h. Middle: same as left, sampling 1000 times a random central spot. Right: same as left, sampling 1000 times a random gene set. C1-2 were significant over both spatial and gene set sampling (p-value < 0.05). **f.** Optogenetic patterning of neural organoids: an embryoid body expressing the PA-Cre/Lox-SHH-GFP system is photostimulated in a restricted area via laser scanning. The resulting organizer, made of SHH-expressing cells, instructs the neighboring cells to form distinct spatial domains of gene expression. **g.** Imaging of DAPI, SHH-expressing cells marked by NeonGreen (GFP for simplicity) and *FOXA2/OLIG2/NKX6-1* in adjacent cryo-sections of neural organoids with laser induction of *SHH* in the north-west pole. Signal in grey scale for each target separately, and merged in green and magenta (right). Scale bar: 100 μm. Experiment was performed in 4 replicates and representative images are shown here. **h.** Molecular Cartography spatial transcriptomic data of control (left) and SHH-induced (right) organoids, with the indicated transcripts colored according to the legend (right). Experiment was performed in 4 replicates per condition and three examples are shown here (one CTR and two SHH). Scale bar: 100 μm. **i.** Distance distribution (in μm) of cells expressing the indicated transcripts from the nearest SHH^+^ cell, in the most left induced organoid of panel h.

To systematically profile spatial gene expression upon local gene perturbations, we established a method to transfer cultured cells on a spatial transcriptomic slide (10X Visium). We adapted hiPSC culture and photo-stimulation to cell culture inserts, which consist of a PET membrane held by a plastic scaffold within a cell culture dish. The membrane can be cut from the scaffold, transferred onto a slide and removed after cell fixation (Fig. 3a, 3c and S4h). We used this system to probe the gene expression response to the induction of *SHH* in the center of the membrane for a time course of 120 hours, with the PA-Cre/Lox system. Since the RNA capture was not homogeneous, yielding vastly different UMI counts across the capture area (Fig. 3d and S4i), we merged the transcript counts for a set of concentric circles, with the inner circle enclosing the photo-stimulated area and the outer ones placed at increasing distances (Fig. 3d). We examined a gene set comprising *SHH* and its targets, and retrieved a peak of expression in the inner part of the membrane for all time points after induction, as compared to randomized controls (Fig. 3e and S4j-l). We note that the raw transcript counts for these transcripts were low (globally, in the range of tens or hundreds), and dominated by *SHH* at the earliest time-point. When considering additional genes in the SHH pathway, we found that the receptor *PTCH1* was strongly upregulated in proximity to *SHH* at 120h, as was, to a lesser extent, its interactor *SMO* (Fig. S4l).

### Optogenetic patterning of neural organoids

*SHH* induction produced a biological response in hiPSCs only detectable with sensitive techniques in short time intervals (by qRT-PCR and to a lesser extent by Visium), or becoming more robust over time as with the *FOXA2* protein becoming detectable 6-7 days after the stimulation. To overcome the constraint of a 2D system which has limited endurance in culture and poor physiological resemblance to a developing tissue, we devised a protocol for producing 3D neural organoids, partially based on previous attempts at mimicking the dorsal-ventral patterning of the caudal part of the neural tube *in vitro* (Zheng et al., 2019). We used laser scanning to induce *SHH* expression in a pole of embryoid bodies grown for 4 days (Fig. 3f and suppl. video 2), then supplemented the medium with retinoic acid for 5 days to induce a posterior fate, and allowed them to grow and differentiate for an additional 7 days (Fig. S5a). We tracked the *SHH* tag fluorescent signal to assess spread and location of SHH-expressing-cells in whole-mount fixed organoids, and noticed that some exhibited significant spread away from the induced pole, likely due to cell divisions and migration. Nevertheless, the organoids retained an overall polarized *SHH* expression, which induced robust and spatially restricted activation of *FOXA2*, whereas non-induced organoids produced neither detectable *SHH* nor *FOXA2* at the protein level (Fig. S5b). We stained consecutive organoid cryosections for *SHH* targets known to be induced in different neural tube domains at increasing distance from the *SHH* source (Ribes and Briscoe, 2009) and observed that *FOXA2* and *OLIG2* established mutually exclusive expression domains resembling *in vivo* neural tube patterning, with *FOXA2* being activated in the proximity of SHH-producing cells and *OLIG2* further away (Fig. 3g). *NKX6-1* expression, instead, encompassed both *FOXA2* and *OLIG2* domains as expected, and was spread across the entire induced organoids (Fig. 3g).

To more comprehensively characterize the spatial patterning activity of *SHH*, we optimized a protocol for performing multiplexed FISH-based spatial transcriptomics on organoids cryosections and imaged a panel of 88 transcripts in parallel, including known *SHH* targets and markers for distinct Dorsal-Ventral (DV) neural tube cell populations. We performed such analysis on four control and four SHH-induced organoids: while control organoids displayed some surface expression of *SHH* and its immediate targets *FOXA1* and *FOXA2*, photostimulated organoids were strongly patterned into distinct spatial gene expression territories (Fig. 3h). Additional markers for all ventral domains (*NKX2-2*, *OLIG1/2*, *NKX6-1/2*, *DBX1/2*) were strongly induced upon *SHH* activation, while exclusively dorsal markers such as *PAX7* and *MSX2* were almost completely depleted (Fig. S5c-e). The spatial distribution of these transcripts resembled that of the neural tube DV axis, consisting of mutually exclusive territories defined by the expression of single or combined markers expressed at defined distances from the *SHH* source (Fig. 3i and S5d). For example, *FOXA1/2*, *NKX2-2* and *OLIG2* expression domains were depleted of *PAX6* and *IRX3*, while *DBX1*, *DBX2*, *OTP* and *OTX2* expression was confined to spatially distinct regions farther away (Fig. S5d). Interestingly, we found that *FOXA1* and *FOXA2* expression pattern differed as *FOXA1* was mostly confined to *SHH*^+^ cells and *FOXA2* was also spread in the immediate vicinity in a non-cell autonomous manner (Fig. 3i). These features are observable in the physical space of single organoid sections (Fig. S5d), as well as in the gene expression space of 43,230 segmented cells from the eight organoids (Fig. S5e).

From these data, we conclude that localized activation of *SHH* signaling can induce spatially restricted patterns of RNA expression in neural organoids, marked by genes known to specify distinct populations of progenitor cells *in vivo* and closely resembling their spatial relationships in the vertebrates’ neural tube.

### Molecular effects of *SHH* and spatial reconstruction of human dorsal-ventral gene expression patterns in neural organoids

From the previous experiments, we assessed that optogenetic activation of SHH in neural organoids induced the expression of known marker genes in well-defined spatial territories. The effects of *SHH* activation on the whole transcriptome, and to what extent these domains resembled those *in vivo* in terms of global gene expression, however, remained unclear.

To address these questions, we performed single-cell RNA sequencing of two pools of neural organoids: control and SHH-induced. After normalization, dimensionality reduction, quality filtering and clustering (Fig. S6a-b), we observed four major cell types: two clearly distinct *SOX2*^+^ neuronal progenitor populations from control and SHH-induced organoids, the latter marked by increased expression of *SHH* and its targets, *DCX*^+^ neurons in similar number from both conditions, and an additional cluster specifically present in SHH-induced organoids and marked by genes encoding extracellular matrix components such as *COL3A1*, *LAMB1* and *GPC3* (Fig. 4a-b and S6a-d). It has been observed that differentiation of oligodendrocyte progenitors can yield a number of *COL3A1*^+^ pericyte-like cells (Marques et al., 2018, Chamling et al., 2021), and our data suggest that *SHH* induction can specifically stimulate this differentiation trajectory.

**Figure 4.**
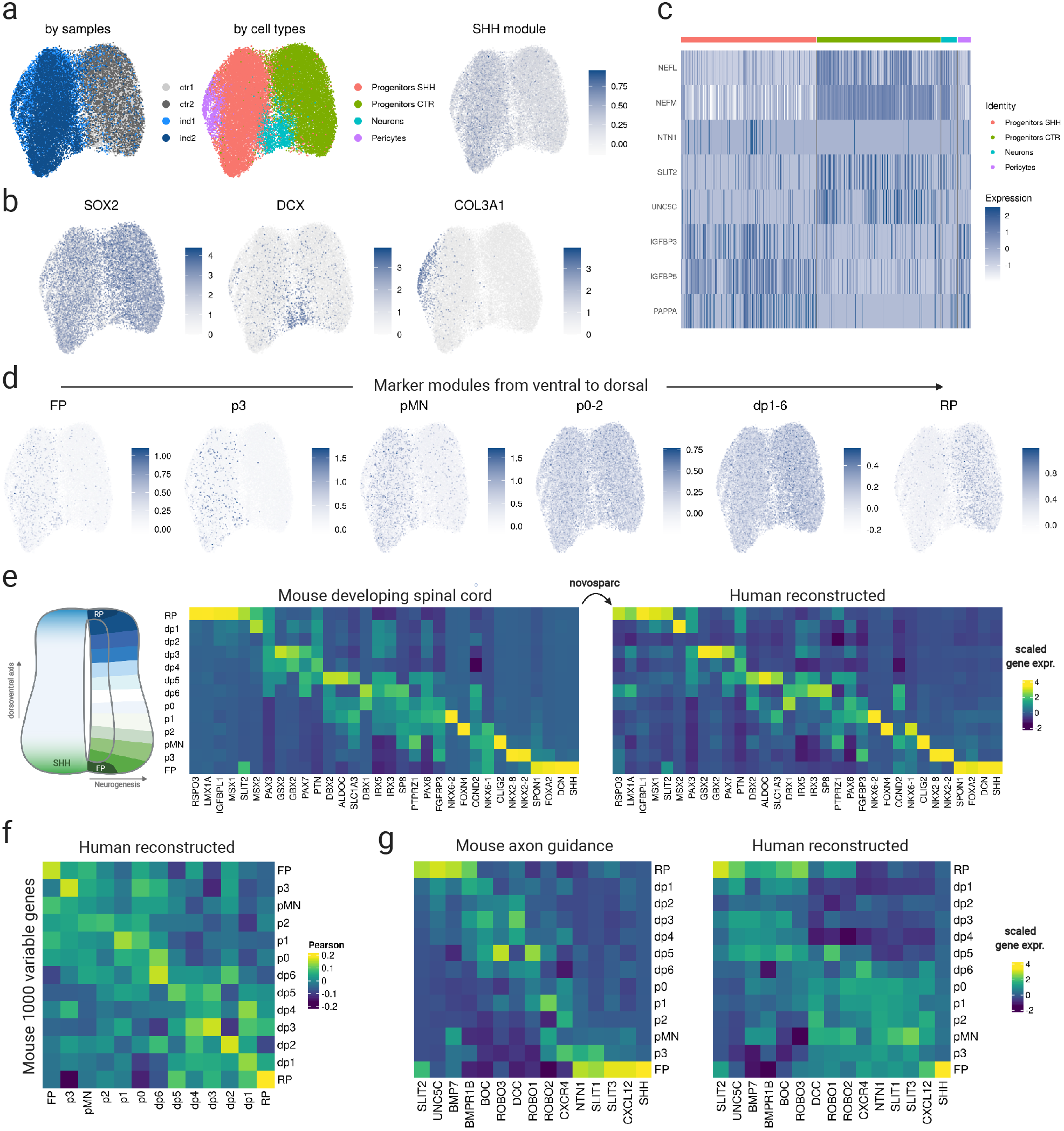
Molecular effects of *SHH* and spatial gene expression patterns in neural organoids. **a.** Sample indication (left), cell types (middle) and *SHH* module score (right) from 2 replicates of control and SHH-induced organoids in UMAP space. **b.** Featureplot of log-normalized expression of markers for main cell types. **c.** Heatmap of log-normalized expression for selected examples of differentially expressed genes. **d.** Featureplot of module scores for markers of dorsal (floor plate, FP) to ventral (roof plate, RP) domains of gene expression of the mouse developing spinal cord. **e.** Left: scaled (z-score) expression of positional marker genes in progenitor cells of the mouse developing spinal cord. Right: reconstructed spatial expression of the same genes in human progenitor cells from control and induced organoids. **f.** Correlation matrix for mouse vs human reconstructed DV gene expression domains, computed on 1000 highly variable genes subsequently filtered for expression in human and mouse datasets. **g.** Scaled (z-score) expression of a set of genes involved in axon guidance in progenitor cells of the mouse developing spinal cord and in the spatial reconstruction of human progenitor cells from control and induced organoids.

Furthermore, by comparing differentially expressed genes between control and induced progenitors (Fig. 4c and S7a), we found genes and gene sets involved in shaping the neuronal cytoskeleton (e.g. neurofilaments *NEFL* and *NEFM*), axon growth and guidance (e.g. *NTN1*, *UNC5C*, *SLIT2*), and regulators of the IGF pathway specifically activated in SHH-induced progenitors (*IGFBP3*, *IGFBP5* and *PAPPA*).

To retrieve the positional identity of the sequenced cells compared to embryonic development, we used the following approach. First, we considered the expression of HOX genes and additional markers of the Anterior Posterior (AP) axis (Philippidou et al., 2013), which indicated that the sequenced cells bear similarity with the posterior part of the hindbrain and the anterior part of the developing spinal cord (Fig. S7b), with the exception of *OTX2*, a midbrain marker which was expressed in a narrow and spatially distinct subset of cells (Fig. S5d).

Second, we compared known positional marker genes along the DV axis of the mouse developing spinal cord (Delile et al., 2019) for its 13 distinct domains, from the floor plate (the most ventral and closest to the *in vivo* SHH organizer), 11 progenitor domains (p3, pMN, p0-2, dp1-6), and the most dorsal roof plate. In concordance with *SHH*’s known role as a ventralizing signal in neural tube patterning, we observed a clear ventralization of cells within the SHH-induced organoids. In comparison, the most dorsal markers of the neural tube were largely confined to cells from non-induced organoids, while markers for intermediate domains were expressed in both conditions (Fig. 4d and S7c), as previously observed by spatial transcriptomics.

Given that marker genes for all 13 domains were expressed in either induced, control, or both organoid conditions, we reasoned that by using merged data from all samples we may be able to reconstruct a global spatial gene expression pattern of a human neural tube DV axis. To do so, we used novosparc (Nitzan et al., 2019), which uses optimal transport to probabilistically place cells in a predefined geometry – in this case a column of 13 DV domains – under the assumption that differences in gene expression are to be locally minimized. By informing novosparc with a set of 32 positional markers retrieved from a mouse developing spinal cord atlas (Delile et al., 2019), we were able to obtain a remarkably similar DV pattern in the human organoids data (Fig. 4e), with cells from induced organoids preferentially assigned to domains from the floor plate to dp6, and cells from control organoids more prevalent from dp5 to the roof plate (Fig. S7d). This was in accordance with the spatial transcriptomics data, which showed that *PAX7* and *MSX2*, expressed dorsally to dp6 *in vivo*, were almost completely depleted in induced organoids (Fig. S4c-d).

When comparing this human reconstruction with mouse *in vivo* data genome-wide, we observed a clear signal along the diagonal of a correlation matrix across the DV domains (Fig. 4f and S7e). However, the correlation coefficients were low in absolute terms, suggesting, as we could expect, that *in vitro* modelling of neural tube development can recapitulate certain gene expression programs observed *in vivo* but presents also substantial differences. Nonetheless, our reconstruction allowed us to examine genes regulated along the *in vitro* DV axis and to validate their expression pattern with *in vivo* data. Interestingly, we observed that the genes encoding the main determinants of axon guidance in the developing spinal cord were strongly regulated along the reconstructed DV axis. This gene set consists of pairs of ligands and receptors (e.g. *SHH-BOC*, *NTN1-DCC*, *ROBOs-SLITs*, *BMP7-BMPR1B*) controlling the establishment of complex neuronal networks across different DV and AP domains of the neural tube (Comer et al., 2019). The spatial expression of these genes was similar to that *in vivo* (Fig. 4g), suggesting that our organoid models could be used for studying important aspects of hindbrain/spinal cord neuronal connectivity *in vitro*.

However, we also observed remarkable differences: for example, the transcription factor *SIM1*, which is expressed ventrally in the mouse developing spinal cord and hypothalamus, was instead enriched dorsally along our reconstructed DV axis, and was in fact downregulated in SHH-induced organoids (Fig. S7f). This observation might result from intrinsic differences between the *in vitro* and *in vivo* conditions, or suggest that *SIM1* expression could be modulated or restricted by *SHH* signaling according to some specific spatiotemporal mechanism in human development that we do not yet understand.

## Discussion

From insects to vertebrates, the key to generate complex body plans from a single cell involves an orchestrated series of patterning events, arising from the definition of spatially restricted gene expression territories during early development. To study these phenomena, we need means to chart gene expression in tissue space with high sensitivity and resolution, and models for reproducing and dissecting the regulatory principles behind them.

The pursuit of imaging RNA/protein expression in tissues dates back to decades ago. *In situ* hybridization and immunostainings have been instrumental to understand the complex interplay of cells in tissues in health, disease and development. In the past few years, spatial investigation of gene expression has made enormous advances, with new technologies for high-throughput spatial transcriptomics (Moses and Pachter, 2021).

Meanwhile, in the field of stem cell-based tissue modelling, the use of three-dimensional organoid culture has gained great interest for its capability to reproduce fundamental aspects of development *in vitro* (Kelley and Pasca, 2022). One missing feature of these technologies is the ability to physically control the spatial patterning of organoid tissues, as it happens during development through the localized expression of morphogens. Controlling the concentration of signaling molecules in the culture media (Zheng et al., 2019) can only generate stochastic asymmetries, and producing assembloids (Cederquist et al., 2019) may have intrinsic limitations in terms of resolution and spatial control.

To overcome these limitations, we applied optogenetic perturbations of gene expression to organoid models of neurodevelopment, and combined these experiments with single-cell and spatial transcriptomic readouts.

We first developed an experimental setup, for which we provide a general blueprint and the necessary tools, which allows the following: 1) choose a gene or a gene set of interest, 2) perturb it (by activation, knock-down, or both) in a specific area of interest of a 2D or 3D cellular model and 3) measure the impact of the perturbation in the transcriptomic and physical space. To this aim, we implemented and optimized light-inducible CRISPRa-based transcription (Nihongaki et al., 2017), Tet-ON transcriptional control (Yamada et al., 2018) and Cre/Lox recombination (De Santis et al., 2021) and combined these with synthetic gene expression cassettes, including a Cas13-based knock-down module, to provide versatile tools of light-inducible gene perturbations.

To imprint gene expression patterns on a cellular “canvas”, we tested different means for spatial stimulations, including photomasks, laser scanning and digital light projection. While fast and comparatively easy, using a photomask in between a light source (e.g. a LED array) and the specimen is limited in resolution and accuracy. We therefore used a laser scanning setup that provides enough resolution, precision and minimum noise to stimulate a single cell. A key limitation of the scanning-based system is that only a single volume of a predefined shape can be stimulated at a time (while, with more sophisticated setups, multiple or complex patterns could be stimulated sequentially). A promising and highly versatile alternative is to employ a DMD setup, which uses a programmable array of micro-mirrors to simultaneously illuminate multiple and complex regions of interest (as for example in Yamada et al., 2018). We engineered such a setup with a digital light processing projector and showed that we can project complex photo-stimulation patterns on cells, inducing gene expression in defined ROIs which can be easily drawn in a graphical user interface.

Spatial gene perturbations may be extremely useful in studying numerous biological processes. For example, overexpression of oncogenes or knock-down of tumor suppressors in a single cell or in defined cell types within organoids may be used as a novel model for tumorigenesis, by examining proliferation and migration of the perturbed cells in the tissue with extreme spatio-temporal control. Another interesting application is the study of developmental processes involving cell-cell interactions. For example, inducing or knocking down signaling molecules or receptors can help us understand the principles and kinetics of cellular interactions in a simplified setup, where the position of sender or receiver cells can be programmed by photo-stimulation (e.g. Rogers et al., 2020).

As a paradigm of this sort of application, we focused on the Sonic Hedgehog pathway. *SHH* is a well-known morphogen produced during vertebrate development by the notochord and the floor plate of the neural tube. It is released in the extracellular space, where it forms a gradient along the dorsal-ventral axis, which specifies the fate of progenitor cells depending on distance from the source of the signal. To imitate this process *in vitro*, we locally induced *SHH* in an artificial organizer and measured its effect on gene expression over time. There have been several studies on *SHH* signaling *in vitro*, by means of treatment with a recombinant protein (e.g. Kutejova et al., 2016), its overexpression (e.g. Li et al., 2018) or fusing a SHH-expressing spheroid with organoids (Cederquist et al., 2019). Very recently, De Santis et al. (2021) used a light-inducible Cre/Lox system to overexpress *SHH* in human induced pluripotent stem cells and showed that this approach can induce strong cell-autonomous and non-autonomous responses. We adopted this approach, as well as a CRISPRa-based system to activate the endogenous *SHH* locus, and combined it with spatial transcriptomics, by establishing a protocol for transferring a cell monolayer to a glass slide suitable for analysis by 10X Visium. We could indeed observe the pathway’s induction and a distance-dependent response, but with limited resolution and sensitivity.

To test the potential of this approach to reproduce tissue patterning in a three-dimensional model of neurodevelopment, we optogenetically induced *SHH* in a restricted group of cells within organoids aimed at mimicking the posterior neural tube development. To quantify *SHH* patterning activity in this context, we optimized single-molecule FISH-based spatial transcriptomics on organoids cryosections for the detection of a custom-designed panel of 88 genes, including general neurodevelopment markers and genes linked to *SHH* biology. Localized *SHH* activation was sufficient to establish distinct neuronal progenitors’ territories, marked by the expression of genes known to specify the dorsoventral identities of the neural tube. For example, we observed that expression of ventral markers *FOXA1/2*, *NKX2-2* and *OLIG1/2* was activated in distinct domains at defined distances from the *SHH* source, where *PAX6* and *IRX3* were instead repressed. The expression of the more dorsal markers *DBX1/2* were restricted to narrow territories located farther away, while *PAX7* and *MSX2* expression was almost completely abolished by *SHH*. In essence, the localized activation of *SHH* signaling was sufficient to reproduce the spatial patterning of the ventral part of the neural tube, from the floor plate to dp5 progenitors, at least according to the expression of well-known positional marker genes along the dorsoventral axis.

Moreover, single-cell RNA-seq data from control and induced organoids revealed that *SHH* could stimulate the differentiation of pericyte-like cells, and strongly induced the expression of the IGF pathway modulators *IGFBP5*, *IGFBP3* and *PAPPA*. *IGFBP5* was proposed to be a *SHH* target in chicken development (Allan et al., 2003) and to regulate the SHH pathway in the cerebellum (Fernandez et al., 2010). *IGFBP3* was proposed to be induced by *SHH* in the fetal prostate (Yu et al., 2009), while *PAPPA*, encoding a protease which specifically cleaves IGFBPs, has not been linked to *SHH* activity to the best of our knowledge. The fact that these three genes appear among the most highly up-regulated genes upon *SHH* activation in neuronal progenitors suggests that they might play a central role in human neurodevelopment and neural tube patterning.

Finally, we used single-cell RNA-seq data to reconstruct human dorsal-ventral gene expression patterns *genome-wide*, by applying an optimal transport-based spatial reconstruction approach, informed by a set of positional marker genes selected by analyzing *in vivo* mouse development data. This reconstruction revealed remarkable similarities among our *in vitro* human model and *in vivo* mouse development – for example, the spatial distribution of genes involved in axon guidance and neuronal connectivity – as well as interesting differences, such the observed repression of the transcription factor *SIM1* by *SHH*.

In conclusion, we believe that our approach of combining optogenetic methods with spatial transcriptomics might prove extremely useful for generating and characterizing new organoid models with complex and controlled spatial patterning modalities, as well as for studying spatio-temporal mechanisms of *SHH* signaling in particular, or gene regulation in general.

## Acknowledgements

We thank Gwendolin Matz and Ruth Alcaraz Pareja for help with cloning and cell culture, Margareta Herzog and Alex Tschernycheff for administration, Salah Ayoub for RNA-sequencing experiments, Sebastian Ehrig for helping with photomasks printing and for casting the PDMS supports for organoids induction, Tamas Sztanka-Tóth, Enes Senel and Nikos Karaiskos for help with Spacemake and Novosparc, Jonathan Froehlich, Giuseppe Macino, Andreas Möglich and Peter Hegemann for useful discussions, Anna Löwa (BIMSB Organoid platform), Sebastian Diecke and Ralph Kuhn (MDC stem cell core facility) for advice on hiPSC culture and transfections, Omar Abudayyeh, Jonathan Gootenberg and Feng Zhang for sharing plasmids and advice regarding Cas13b, Riccardo De Santis for sharing the Cre/Lox plasmids, advice regarding the Cre experiments and feedback on this manuscript, Yuta Nihongaki for advice regarding CPTS/split-CPTS optimization, Itaru Imayoshi and Mayumi Yamada for sharing the PA-TetON plasmids, Heiko Lickert for sharing the XM001 hiPSC line, the whole N. Rajewsky lab for discussions, support and constructive feedback, David Koppstein and Christine Kocks for reading the manuscript, Tatiana Borodina, Caroline Braeuning, Thomas Conrad, Daniele Yumi Sunaga-Franze, Kerim Secener, Jeannine Wilde (BIMSB sequencing facility) for sequencing, Visium and FACS experiments. We thank the Resolve Biosciences team for performing the Molecular Cartography assays and data processing. All illustrations were created with BioRender.com.

## Author contributions

NR and IL conceived the original idea. NR supervised the whole project, giving input about experiments and analyses. IL wrote the first manuscript draft with input and feedback from all authors, NR revised and finalized the manuscript. IL designed and supervised all experiments with input from NR, performed all Cas13 experiments in HEK cells, cloned most of the plasmids, performed RNA extractions and qPCRs, performed most of the inductions with the LED board, the confocal and the DMD setups, analyzed qPCR, imaging, bulk and single-cell RNA-seq, Visium and Molecular Cartography data. LE performed all experiments in hiPSCs, RNA extractions, qPCRs, immunostainings and optimized the Visium protocol from 2D culture. AB optimized the Visium protocol for 2D culture with LE and produced all Visium libraries. CCJ optimized and performed live cell imaging of GFP and RFP in HEK cells. TH generated the PA-Cre/Lox HEK cell lines, cloned some of the plasmids, performed RNA extractions and qPCRs and assisted LE with immunostainings. GM performed the proteomics experiment and analyses, under the supervision of SK. AOM optimized and performed CasRx inductions with the confocal microscope setup with IL. ARW supervised the work done with hiPSCs, performed cell and organoid culture, produced single-cell RNA-seq libraries and the organoids cryosections for spatial transcriptomic analysis. AW constructed the DMD setup, wrote the code for controlling spatial stimulations and optimized CasRx and SHH inductions with this setup. RW contributed at the initial stage of the project designing, producing and programming the LED board for whole-well stimulations and assisting IL with the first induction experiments. AZ developed the protocol for generating neural organoids, performed SHH inductions with IL, immunostainings and imaging, and produced single-cell RNA-seq libraries, under the supervision of RZ.

## Data availability

Sequencing data have been submitted to GEO and will be made available upon revision of this manuscript.

## Methods

### Cell culture, transfections and cell lines generation

HEK293T cells were cultured in DMEM (high-glucose, with glutamax and pyruvate, Thermo Fischer #11360872) supplemented with 10% Tet-free FBS (PAN biotech., #P30-3602) in absence of antibiotics at 37°C with 5% CO_2_. They were split every 2/3 days with 0.05% trypsin in 10 cm cell culture dishes. For transfection experiments shown in figure S3, 30,000 cells were seeded the evening before transfection in 70 μl medium on white 96-well, clear-bottom plates (Corning #3610). The morning after, a mix comprised of 12 μl Optimem (Thermo Fisher, #31985062), 25 ng Luciferase-encoding plasmid, 150 ng guide RNA-encoding plasmid and 150 ng Cas13-encoding plasmid or 2 times 100 ng of each Split-Cas13-encoding plasmid was mixed with 25 μl Optimem and 0.5 μl Lipofectamine 2000 (Thermo Fisher #11668019), incubated at room temperature for 10 minuets and then pipetted onto the cells. Light stimulation was started 6 hours post-transfection and luciferase assay was performed in the same plate 24 hours post-induction by removing 50 μl medium, adding 75 μl Luciferase assay buffer (Promega Dual-Luciferase Reporter Assay System, #E1910), incubating 10 minutes at room temperature, then reading Firefly luciferase, then 75 μl Stop&Glo buffer, incubating 10 minutes at room temperature, then reading Renilla luciferase. Plate readings were performed in a Tecan M200 infinite Pro plate reader with two-seconds integration for luciferase measurement.

For transfection experiments shown in figures 1, 2, S1 and S2, 25,000 cells were seeded the evening before transfection in 70 μl medium on black 96-well, clear-bottom plates (Corning #3904). The morning after, a transfection mix comprised of 25 μl Optimem, 0.4 μl P3000 and 100-300 ng plasmid DNA was pooled with 25 μl Optimem and 0.3 μl Lipofectamine 3000 (Thermo Fisher #L3000001), incubated for 10 minutes at room temperature and then pipetted onto the cells. Light stimulation was started 6 hours post-transfection and live-cell imaging was performed at 24 hours post-induction unless differently indicated (e.g. for the time-course in figure 1).

Transfections for the RNA-seq and proteomics experiments shown in figure S2 were performed in 6-well plates with 1 million HEK293T cells per well. The morning after seeding, a mix comprised of 250 μl Optimem, 800 ng ePB Puro TT RFP plasmid, 1000 ng guide RNA-encoding plasmid and 1000 ng Cas13-encoding plasmid, 8 μl p3000 reagent and 2 μl doxycycline (1mg/ml) was mixed with 250 μl Optimem and 6 μl Lipofectamine 3000, incubated at room temperature for 10 minutes and then pipetted onto the cells. Cells were harvested 36h post-transfection for RNA extraction with home-made trizol or for protein purification, as described later in the RNA-seq and proteomics sections.

Transfections for generating the Cre/Lox CasRx line were performed in 12-well plates seeded with 100,000 HEK293T cells and the day after transfected with 250 ng ePB-PA-Cre plasmid, 500 ng LoxP-CasRx plasmid, 125 ng hyperactive transposase plasmid, 100 μl Optimem and 2.5 μl p3000 reagent, mixed with 100 μl Optimem and 1.5 μl Lipofectamine 3000 reagent, incubated for 10 min at RT and then pipetted onto the cells. After 3 days post-tranfection, cells were split into two wells of a new 6-well plate and selected with puromycin (1 μg/ml) and blasticidin (5 μg/ml) for a week. RFP^+^/GFP^-^ cells were further FACS-sorted for increasing the purity of the population.

HiPSCs (XM001 line, Wang et al., 2018) were cultured in E8 flex media with supplement (Thermo Fisher #A2858501) at 37°C in hypoxia (5% O2) conditions, passaged every 3-4 days with accutase, and seeded on Geltrex-(Gibco #A1413302) coated plates. To promote their attachment to the plates, cells were kept in E8 flex media supplemented with 10 μM Rho-associated protein kinase (ROCK) inhibitor. Media was changed to E8 flex without ROCK inhibitor within the next 24h from plating.

The PA-TetON CasRx cell line was produced with two lentiviruses produced in HEK293T cells with the PA-TetON plasmid (Yamada et al., 2018) and the TRE-CasRx plasmid, two packaging plasmids (Addgene #8454 and 8455). 50,000 HEK293T were transduced with a MOI of 10 of each lentivirus in a 24-well format and then selected for blasticidin expression (5 μg/ml) for one week.

HiPSCs transfections were conducted using Lipofectamine Stem (Thermo Fisher #STEM00001). For transfections experiments regarding the timecourse of SHH activation shown in Figures 3 and S4, hiPSCs colonies at 70-80% confluency were dissociated to single cells with accutase. 250,000 cells were then resuspended in 100ul of E8 flex with 10 μM ROCK inhibitor and seeded on black 96-well plates, previously coated with Geltrex. After 2-3 h, when cells got attached to the wells, media was changed with 50 μl of E8 flex with ROCK inhibitor. The transfection was performed as follows: for one sample, 1.2 μl of Lipofectamine were mixed with 25 μl of Optimem and 500 ng of total plasmids were diluted in 25 μl of Optimem. Diluted Lipofectamine and diluted plasmids were then combined at a ratio 1:1, incubated at room temperature for 10 minutes and 50 μl pipetted drop by drop on top of the cells. Transfection efficiency was assessed by including a control plasmid encoding for constitutive GFP (pmax-GFP, Lonza, #V4YP-1A24). Media was changed within 7-8 hours after transfection to E8 flex and, prior to light induction, gradually replaced by neural induction media “COM1”, whose composition is the following: DMEM-F12 (Thermo Fisher #11320033), N2 supplement (Thermo Fisher #17502048), Neurobasal (Thermo Fisher #21103049), B27-vitamin A (Thermo Fisher #12587010), 1X Penicillin/Streptomycin, Glutamax (Thermo Fisher #35050061), 2-mercaptoethanol, vitamin C, CDLC (chemically defined lipid concentrate) and insulin. The reason of this media switch was due to the fact that E8 flex contains basic fibroblast growth factor (FGF2), which is a strong inhibitor of the SHH signalling pathway (Fogarty et al., 2007). Light stimulation was started within 15 hours post-transfection, using blue LED array. Cells were harvested after each time point of light stimulation and lysed with 100 μl home-made Trizol. RNA was then extracted with the Zymo RNA extraction kit (Zymo #R2051). cDNA preparation and qRT PCR were performed as described in paragraph “RNA extraction, qRT-PCRs”.

The plasmids encoding the SCPTS 2.0 systems were generated as previously described with standard molecular cloning approach (Nihongaki et al., 2017), and with the aim to produce stable iPSC lines, the modules in these plasmids (split Cas9, MCP, sgRNA cassette) were PCR-amplified and recloned into 2 lentiviral plasmids (pLKO.1 neo, Addgene #13425; pLJM-EGFP, Addgene #19319) and, from the latter, they were subcloned into two PiggyBac transposons vectors. With the SCPTS2.0 system we did not succeed in generating stable lines, neither by lentiviral infections nor by transposase/PiggyBac strategy: in both cases, we experienced a full cell mortality, perhaps due to some forms of toxicity of the system. On the other hand, we managed to generate stable iPSCs Cre/Lox SHH and Cre/Lox CasRx lines by transfecting hiPSCS with PiggyBac vectors and the transposase enzyme. In this regard, 400,000 hiPSCs were plated in a Geltrex-coated well of a 12 well-format plate and transfected with lipofectamine. Herein, 400ng of each transposon-one carrying the “TRE-CRE split1 nMag - T2A/P2A-pMag CRE split2” and the other carrying the “CAG-loxP-RFP-loxP-CasRx or SHH cassettes”-were combined with 200 ng of hyperactive transposase and diluted in 72 μl of Optimem. Diluted DNA was then mixed with lipofectamine (3 μl), also diluted with Optimem (72 μl) and incubated at RT for 10 minutes. More specific details of the transfection protocol are outlined above. Media was changed to E8 flex after 5h from transfection. Cells were let recover for 4 days and antibiotics selection was then started, immediately after splitting hiPSCS: 1 μg/ml of puromycin and 2 μg/ml of blasticidin were added to E8 flex medium. Cells were kept under antibiotics selection for 10 days. As a readout of successful integration of the transposons cassettes, RFP signal was checked.

### Organoids differentiation

HiPSCs were cultured in E8 flex medium (Gibco #A14133-01) with medium replacement every other day until 80% confluency, in the dark. The differentiation protocol was adapted from Zheng et al. (2019). HiPSC colonies were rapidly washed with PBS (Pan Biotech P0436500) and then incubated with accutase (Sigma, #A6964-100ML) for 4 minutes at 37°C. Cells were collected, centrifuged for 3 minutes at 300 g and resuspended in E8 flex medium containing 10 μM Y27632 ROCK inhibitor (VWR, #688000-5). Cells were counted and plated at a density of 500 cells per well in a 96-well ultra-low attachment U-bottom plate (Corning, #CLS7007). On the following day, the medium was replaced with fresh N2-B27 medium (50% Neurobasal (Gibco #A3582901), 50% DMEM/F12 (Gibco, #11320074), 1x N2 (Gibco #17504044), 1x B27 (Gibco, #17504044), 1x MEM non-essential amino acids (Sigma; M7145-100ML), 1x Glutamax (Gibco, #35050038), 0.1 μM β-mercaptoethanol (Merck Millipore #8057400005) supplemented with 2% Geltrex (Gibco, #A1413301), 10 μM TGF-β pathway inhibitor (SB431542, Stem cell technology, #72234) and) 0.1 μM BMP inhibitor (LDN193189, Stem cell technology, #72147). Medium was exchanged daily. From day 4 on, organoids were cultured in 35 mm dishes, medium was additionally supplemented with 1 μM retinoic acid (RA, Sigma, #R2625) until day 8. At day 9, RA was removed, organoids were further cultured in N2B27 medium supplemented with LDN and SB until day 16. For organoids optogenetic stimulations, the same protocol as before was used, but the medium was supplemented with 1 μg/ml doxycycline to activate PA-Cre expression at day 3. On the next day, four organoids at once were embedded in a drop of Geltrex on a glass bottom dish (WillCo-dish, #GWSB3522), incubated for 15 min at 37°C and covered with warm N2B27 medium (supplemented with SB, LDN and RA). To induce *SHH* expression in a restricted pole of the organoids, the laser scanning setup was chosen (Leica Sp8 SMD, see suppl. video 2): a small square ROI of ca. 100-400 μM was selected depending on how the four organoids were positioned with respect to each other, induced for two times 5 seconds at 100Hz with 1% laser power set at 480 nm, every 30 seconds overnight (16 hours). After induction, organoids were retrieved with a pipette and cultured individually until day 16 in an ultra-low attachment 24-well plate (Corning #CLS3473). Control organoids were not induced. Media was exchanged daily with fresh N2B27 supplemented with SB, LDN, 2% Geltrex and RA until day 8. From day 9 onwards, RA was removed as described before.

### LED board construction and stimulation experiments

For experimental convenience, we decided to build a custom circuit board with 96 blue LEDs that align with the used 96 well-plates. To control illumination patterns for each well individually we opted to wire each LED to a dedicated output line of a constant current LED driver chip (MAX6969). Optimizing for brightness at low supply currents to minimize excess heat, we decided on the Cree XLamp MLESBL with a documented center wavelength at 485nm and a reported luminous flux of 13.9 lm at 50 mA. We soldered the 96 blue LEDs onto a custom aluminum PCB. The LED PCB serves as a heat sink and is exposed to the incubator environment. A dark PVC hole mask reduces light spill. The assembly is encased in an acrylic frame and the seams sealed with neutrally curing silicone. Two cables leave the case: one to control the shift registers of the driver chips with a micro-controller outside of the incubator, and another to power the LEDs (7-9 V) and the logic chips (5V). We used the serial interface of a micro-controller (Atmel AVR ATmega32) to periodically update the shift registers of the LED drivers according to the desired patterns and to control the output latches. We opted for a control frequency of 1Hz and specified the illumination patterns with a simple domain-specific language supporting four instructions: turn on the LED for up to 127 seconds (0×00...0×7E), turn off the LED for up to 127 seconds (0×80...0×FE), repeat the pattern (0×7F), and halt (0×FF). The code and the schematics for the LED board and the LED drivers are available at https://github.com/BIMSBbioinfo/casled. Stimulations were performed with a 5-seconds on, 20-seconds off pattern repeated over the desired time interval (usually 24 hours), with the cell culture plate placed directly on top of the LED board. To avoid heating, input voltage was set at 7.6V for most experiments (below the optimal value for the LEDs used) and temperature of the medium in a lit 96-well plate was checked in a preliminary test with a thermocouple.

### Experimental setup for parallel optogenetic stimulation (DMD setup)

Illumination from a DMD-based projector (DLP LightCrafter 4500, Texas Instruments, modified for on-axis projection by EKB technologies) was coupled to the rear port of an Observer.Z1 microscope (Zeiss), through a unity magnification relay (2x AC254-125-A-ML, Thorlabs) with an OD 2.0 neutral density filter (NE20A-A, Thorlabs). For optical stimulation, illumination from the blue (470 nm) LED of the LightCrafter passed through a GFP filter set (ET-GFP, Chroma, Bellow Falls, VT, USA) and projected to the sample with a 10× Plan Apo objective. For imaging of RFP, the green (530nm) LED was used together with a CY3 filter set (ET-CY3/TRITC, Chroma). Projector / camera pixel mapping and subsequent control of illumination patterns was performed using the projector plugin for Micromanager 2.0 gamma (EdelStein et al., 2014). Illumination intensity was controlled using DLP LightCrafter 4500 Control Software (v3.1.0, Texas Instruments). Emission was detected by a back illuminated sCMOS (PrimeBSI, Teledyne Photometrics). For optogenetic stimulation, samples were illuminated with 470 nm excitation at a power density of 4.7 μW/cm^2^ in user defined regions of interested (ROIs) for 20 seconds. After stimulation, full field of view RFP images were acquired. This was repeated every minute for 16 hours using a custom written Beanshell script in Micromanager. Environmental control during long term time-lapse imaging was achieved with the Incubator XLmulti S chamber and temperature/CO2 controllers (PeCon, Germany).

### Laser scanning setup for single-cell stimulations

Scanning-based optogenetic stimulation experiments were conducted using a confocal microscope Leica TCS SP8 (Leica Microsystems) equipped with an environmental (CO_2_ and temperature) control system. Imaging and stimulation were performed using a 10× Plan APO objective and a white light laser tuned to 488 nm at 1% laser power. Scanning-based stimulation of 100 x 100 μm ROI containing a single cell, was performed at 100 Hz unidirectional scan speed. Two sequential scans were performed resulting in 10 seconds of total exposure. The stimulation protocol was repeated every 30 seconds for 16 to 20 hours. The scanning-based stimulation setup mimicked the previous LED stimulation pattern, although scanning time was set to 10 seconds, instead of 5 seconds LED illumination, in order to correct for the off-sample scan time.

### RNA extraction, qRT-PCRs

For the experiments shown in figures 2 and 4, RNA extraction was performed as follows. Cells were harvested by removing medium from 96-well plates, adding 100 μl home-made Trizol directly onto the cells while keeping the plate on ice, then pipetting up and down a few times and transferring the lysate into a new 1.5 ml tube. Lysates from two or three wells were pooled in each replicate, then RNA was extracted with the Zymo Directzol RNA miniprep kit (Zymo, #R2051), including DNase I digestion. cDNA was synthesized using 100-200 ng RNA with the Maxima H minus RT (Thermo Fisher, #EP0751) according to the manufacturer protocol and using random hexamers for priming. 5 ng of diluted cDNA were used per qPCR reaction using ROX-supplemented Biozym SYBR green mastermix (Biozym, #331416S) and 0.5 μM forward and reverse primers. qPCR reactions were performed in a AB 7500 machine with the following cycling conditions: 95°C for 10 minutes, then 40 cycles of 95°C for 15 seconds and 60°C for 1 minute with fluorescence reading, and final melting curve step.

### LC-MS proteomics

Cells were transfected and treated as described in the dedicated section. They were then checked by fluorescence after 36h for RFP knock-down and processed for proteomic analysis as follows. Cells were resuspended in 350 μl of Urea buffer (8 M Urea, 100 mM Tris-HCl, pH 8.2). Cells were lysed on a Bioruptor sonicator (Diagenode), using 10 cycles of sonication (45 sec ON, 15 sec OFF). Protein concentration was determined by bicinchoninic acid colorimetric assay (Pierce) and a 100 μg aliquot of each protein sample was reduced with 10 mM DTT for 45 minutes at 30 °C and alkylated with 100 mM iodoacetamide for 25 minutes at 25 °C. Proteins were digested using Lys-C (Wako, 1:40, w/w, overnight under gentle shaking at 30°C) and modified trypsin (Promega, 1:60, w/w, 4 hours under rotation at 30°C). Lys-C digestions product were diluted four times with 50 mM ammonium bicarbonate before the tryptic digestion, which was stopped through acidification with 5 μl of trifluoroacetic acid (Merck). Fifteen μg of each resulting peptide mixture were then desalted on Stage Tip (Rappsilber et.al., 2007), the eluates dried and reconstituted to 15 μL in 0.5% acetic acid. For all the samples, 5 microliters were injected on a LC-MS/MS system (EASY-nLC 1200 coupled to Q Exactive HF, Thermo), using a 240 minutes gradient ranging from 2% to 50% of solvent B (80% acetonitrile, 0.1% formic acid; solvent A=0.1% formic acid in water). Each sample was analyzed in duplicate. For the chromatographic separation 30 cm long capillary (75 μm inner diameter) was packed with 1.9 μm C18 beads (Reprosil-AQ, Dr. Maisch HPLC). On one end of the capillary nanospray tip was generated using a laser puller, allowing fretless packing (P-2000 Laser Based Micropipette Puller, Sutter Instruments). The nanospray source was operated with a spray voltage of 2.0 kV and an ion transfer tube temperature of 260 °C. Data were acquired in data dependent mode, with one survey MS scan in the Orbitrap mass analyzer (120,000 resolution at 200 m/z) followed by up to 10 MS/MS scans (30,000 resolution at 200 m/z) on the most intense ions. Normalized collision energy was set to 26, Once selected for fragmentation, ions were excluded from further selection for 30 s, in order to increase new sequencing events. Proteomics data processing and analysis Raw data were analyzed using the MaxQuant proteomics pipeline (v1.6.10.43) and the built in the Andromeda search engine (Cox et al., 2011) with the Uniprot Human database. Carbamidomethylation of cysteines was chosen as fixed modification, oxidation of methionine and acetylation of N-terminus were chosen as variable modifications. The search engine peptide assignments were filtered at 1% FDR and the feature match between runs was enabled. For protein quantification LFQ intensities calculated by MaxQuant were used (Cox et al., 2014). The minimum LFQ ratio count was set to 2 and a MS/MS spectrum was always required for LFQ comparison of the precursor ion intensities. Data quality was inspected using the in-house developed tool PTXQC (Bielow et al, 2016). After removing reverse and contaminants hits, LFQ intensities were log2 transformed and proteins with less than four valid values in each condition were filtered out. Proteins with differential expression between conditions were test with Student’s ttest with Benjamini-Hochberg FDR set at 0.05. Processed data are available in the supplementary table 1.

### Plasmids

Supplementary table 2 contains the name, description and information on availability on the plasmids used in this study.

### Bulk RNA sequencing

Cells were treated exactly as for the proteomics experiment and as described in the dedicated section, in two additional replicates per condition. RNA was extracted with home-made Trizol by organic phase separation and RNA precipitation. Total RNA-seq libraries were performed as follows: 1 μg of total RNA per sample was first depleted of ribosomal RNA using the RiboCop rRNA Depletion Kit (Lexogen, #144) according to the manufacturer’s instruction. The rRNA-depleted samples were then processed with the TruSeq mRNA stranded kit from Illumina. Libraries were then sequenced on a Nextseq with 1×76 cycles. Fastq data were generated with the bcl2fastq program and fed to the PiGx analysis pipeline (Wurmus et al., 2018), which was used with default settings with a custom reference GRCh38 human genome supplemented with two extra chromosomes carrying the CasRx-T2A-GFP cassette and the TagRFP cassette, and a custom annotation made of the Gencode v34 human annotation supplemented with two extra entries for the CasRx-T2A-GFP and the TagRFP genes. For further analyses, we used the STAR/Deseq2 PiGx output.

### Live cell imaging for GFP and RFP quantification

After 6, 12, 25 or 50 hours of light induction or the respective dark controls, images for GFP and RFP were acquired on an inverted Nikon Ti-E microscope with a 4x NA1.4 objective and Andor iXON Ultra DU-888 camera; Z stacks had 1.5 μm spacing over a 40 μm range. GFP: 300ms exposure; Sola 50% on 6-12h, 12% on 25-50h. RFP: 100ms exposure; Sola 20%. All these images were taken with live cells in black 96-well plates. Z-stacks were used for max intensity projection within imageJ, and the projection were used for signal quantification with a macro running the imageJ *Subtract Background* plugin with a rolling ball radius of 50, and then the *Measure* function for signal intensity. This quantification assumes that all wells contain on average the same number of cells, which were seeded in the beginning of the experiment. For some of the wells, we noticed a pipetting artifact on a side, producing an area devoid of cells. We manually selected a ROI which excluded this area for all wells, and we applied this ROI before running the signal measurement macro. This experiment was performed blindly: IL transfected the cells and performed the light stimulation, then CCJ performed the imaging without knowing the samples labels, then IL ran the macros and reassigned the original labels to the well names.

### Immunofluorescence of hiPSCs and organoids

HiPSCs or organoids were rapidly washed in cold PBS and fixed in 4% PFA for 10 minutes in a multi-well plate with agitation. For whole-mount imaging, permeabilization and blocking were performed for 1h at room temperature in PBS solution containing 0.1% Triton-X, 0.2% BSA and 4% normal donkey serum. Organoids were subsequently incubated with primary antibodies overnight at 4°C in blocking solution (PBS supplemented with 0.2% BSA and 4% normal donkey serum). The following primary antibodies were used in immunostaining: Anti-FOXA2 (R&D systems, #AF2400; 1:100), Anti-OLIG2 (Sigma, #HPA003254-100UL; 1:1000), Anti-NKX6.1 (Sigma, #HPA036774-100UL; 1:500). On the following day, hiPSCs/organoids were washed 3 times for 10 minutes, with agitation, with washing solution (PBS supplemented with 0.1% Triton-X, 0.2% BSA). Secondary antibodies and DAPI (Sigma, #D9542) were then incubated at room temperature for 1h in blocking solution. The following secondary antibodies were used at 1:1000 dilution in blocking solution: Alexa Fluor 647 anti-Rabbit (Thermo Fisher, #A21244), Alexa Fluor 647 anti-Goat (Thermo Fisher, #A21447), depending on the primary antibody. Samples were then washed again 3 times for 10 minutes, with agitation, in washing solution. For mounting, the organoids were placed in the center of a slide, washing solution was carefully removed and one drop of Prolong Gold antifade reagent (Thermo Fisher, #P36930) was placed on top of each organoid. A coverslip was placed on top and the slides were allowed to dry at room temperature overnight in the dark. For hiPSCs, the mounting media was added directly in the cell culture plates were cells were seeded (Thermo Fisher, #P10144).

For organoids slices: after fixation, organoids were allowed to settle in 1 mL 40% sucrose solution overnight at 4°C. On the following day, they were embedded in 13%/10% gelatin/sucrose solution and positioned inside an embedding mold (Sakura #4566), rapidly moved to dry ice to freeze and then placed at −80°C for storage. Blocks were removed from −80°C and allowed to warm inside the cryostat to sectioning temperature (−20°C) for 15 minutes. Sectioning was performed using a cryostat (Thermo Fisher Cryostar NX70) and set to produce 10 μm-thick slices. Cut sections were collected on slides (Thermo Fisher,#J1800AMNZ) and stored at −80°C for long-term. To perform immunostaining, slides were allowed to warm to room temperature for 10 minutes, incubated for 5 minutes with 37°C PBS to remove embedding solution. Permeabilization and blocking, as well as incubation with primary and secondary antibodies, were done as described above for whole-mount organoids.

Images were acquired using a confocal microscope (Leica TCS SP8) using a 10X dry or a 20× immersion objective. Z-stacks and final images were processed using Fiji-ImageJ, to produce maximal intensity projections and to subtract background.

### Spatial transcriptomics: VISIUM experiments and analysis

PET membranes (Millipore Millicell Hanging Cell Culture Insert, PET 3 μm, 24-well, #MCSP24H48) were positioned in glass bottom black 24-well plate (Greiner Bio-one, #662892), after cutting away the plastic holders, hence making the membrane touch the bottom of the well with no gaps in-between (this step was performed to ensure no light scattering or diffusion). Circular black photomasks were sticked underneath the bottom of the plate. Membranes were coated with 100 μl of cold Geltrex. iPSCs were splitted to single cells, as described above, and 275,000 cells resuspended in 100ul were cultured on coated membranes generating a stable monolayer. Additional warm E8 media (300 μl) was pipetted around the plastic scaffold. Plates were incubated at 37°C for 2/3 hours until cells attachment. Samples were prepared in duplicates with the intent to perform control quantifications for each of those prior to the final Visium experiment. Plates were kept wrapped in aluminum foil to avoid light exposure. Before starting with the first 24h of light induction, media was changed to ½ E8 flex + ½ COM1 and 1 μg/ml of doxycycline.

The plate with the cells to be induced was covered on top by a black velvet lid and positioned onto the blue LED plate. The control sample (0h) was kept in the dark during the whole time course. Media was changed to ¼ E8 flex and ¾ COM1 between 24h and 48h of induction. Finished the time course, the 4 samples were transferred to a Visium Spatial Gene expression slide (10X Genomics) as follows: the plastic structure that surrounds the membrane was carefully held with tweezers and turned upside down to get rid of the media; membranes were delicately washed twice with 100 μl of PBS and, by using a scalpel, delicately isolated from the plastic device. By using tweezers, the membranes were then slowly sticked onto a Visium Spatial Gene Expression slide with cells facing on it. The more the membrane was kept flat, the more efficient the cells transfer. The Visium Spatial Gene Expression protocol was followed, according to manufacturer instructions (10X Genomics.).

Fastq files were first processed for retrieving transcript counts with positional information with the spaceranger software (10X Genomics, v. 1.2.0). The output of spaceranger was loaded into Seurat (v. 4.0) within RStudio with R 4.0.4, and each sample was subsetted into 7 concentric circles with the center being set according to the stimulation pattern observed by fluorescent microscopy (and after checking that different radii would yield stable results in the samples with SHH induction, subsetting from 5 to 10 concentric circles and finally settling for an intermediate size). The central spots selected for each sample from 0 to 120h had the following barcodes: CACATGATTCAGCAAC, CAATTTCGTATAAGGG, CAATTTCGTATAAGGG, GGAGGGCTTGGTTGGC (the latter north-west from the physical center as the induction was not centered). At this point, concentric circles were drawn by taking all spots with a distance from the center < 500, 775, 1050, 1325, 1600, 1825 and > 1825 for the c1-c7 areas. Within these subsets of spots, the transcript counts for a *SHH* gene set comprising *SHH*, *NKX6-1*, *NKX6-2*, *NKX2-2, NKX2-1*, *FOXA2*, *FOXG1* and *OLIG2* were added and normalized for the total transcript counts of each subset, and then further normalized by the mean of the counts for all spatial subsets c1-c7. As controls, we either randomized the genes in the gene set 1000 times, or the center spot 1000 times, and then computed an exact p-value for each subset gene set enrichment testing the hypothesis of the enrichment being larger than the random control. The signal was stable with varying binning sizes (from 6 to 9) and over cumulative analysis per single capture spot.

### Spatial transcriptomics: Molecular Cartography experiments and analysis

Control and SHH-induced organoids at 14 days post-induction were washed twice in PBS, submerged for 2 hours in PaxGene fixation reagent at room temperature (Qiagen #765312), kept overnight at 4°C in PaxGene stabilization reagent, then soaked in a 40% m/v sucrose/PBS solution for 30 minutes, OCT-embedded, snap-frozed and cut in 10 μm-thick cryosections which were placed directly on a glass slide provided by Resolve Biosciences for Molecular Cartography analysis. Slides were shipped to Resolve Biosciences, where they were processed for multiplexed single-molecule fluorescent *in situ* hybridization of a panel of 88 transcripts of interest, including those showed in Fig. 3 and S5 and additional neuronal markers and housekeepers.

Images and transcript quantification data provided by Resolve were processed using Fiji-imageJ and the Polylux v1.9.0 plugin for transcripts visualization over a binarized grey mask of the processed organoids (Fig. 3h and S5d), or imported in R and processed with Seurat v. 4.0.5 on R v. 4.0 for performing log-normalization, producing the heatmap in Fig. S5c with the DoMultiBarHeatmap() package (https://github.com/elliefewings/DoMultiBarHeatmap), and the dimensionality reduction plots (UMAP performed on 12 PCs) in Fig. S5d.

For computing the distance distribution shown in Fig. 3i, the cell segmentation ROIs provided by Resolve were imported in imageJ where they were used for computing their centroids coordinates. The distance of the centroid of cells with at least 5 counts of each transcript of interest was computed from than of each cell with at least 5 *SHH* counts, and only the distances with the nearest SHH^+^ cell were kept for further analysis.

### Single-cell RNA sequencing

Organoids were dissociated using Accutase, followed by washing with growth medium and filtration through 40 μm cell strainer. Cells were then pelleted and resuspended in PBS supplemented with 0.01% BSA. Cells were counted with a hemocytometer and diluted to a suspension of ~ 300 cells/μl. Cells were encapsulated together with barcoded microparticles (Macosko-2011-10 V+, ChemGenes) using the Dolomite Bio Nadia instrument, using the standard manufacturer’s dropseq-based scRNA-seq protocol. Droplets were broken immediately after collection and cDNA libraries generated as previously described (Wyler et al., 2021). First strand cDNA was amplified by equally distributing beads from one run to 24 Smart PCR reactions (50 μl volume; 4 + 9 to 11 cycles). 20 μl fractions of each PCR reaction were pooled, then double-purified with 0.6 volumes of AMPure XP beads. Amplified cDNA libraries were assessed and quantified on a BioAnalyzer High Sensitivity Chip (Agilent) and the Qubit dsDNA HS Assay. Nine-hundred pg of each cDNA library was fragmented, amplified (13 cycles) and indexed for sequencing with the Nextera XT v2 DNA sample preparation kit (Illumina) using custom primers enabling 3’-targeted amplification. The libraries were purified with AMPure XP Beads, quantified and sequenced on Illumina sequencers (first run: concentration 2pM; Hiseq 3000/4000 SBS kit (150 cycles) paired-end mode; read 1=75 using the custom primer Read1custseqB (8bp) read 2=75. Second run: concentration 2 pM; NextSeq 500/550 High Output v2 kit (75 cycles) in paired-end mode; read 1 = 21 bp using the custom primer Read1CustSeqB (8 bp) read 2 = 63 bp).

Data were produced and demultiplexed by the BIMSB genomics platform, and fastq files were used as input to the Spacemake pipeline (Sztanka-Tóth et al., 2021; https://github.com/rajewsky-lab/spacemake), used with default parameters in scRNAseq run mode and with a min. 500 UMI / max. 10,000 cells cutoffs, to generate a gene expression matrix for each of the fastq file. Since two sequencing runs were performed for each sample, Spacemake was used in merge mode to create a merged gene expression matrix for each of the 4 samples. Such matrices were then used as input for Seurat v. 4.0.5 on R v. 4.0 to create a merged Seurat object. The Seurat object was filtered for cells with > 800 UMIs and < 5% mitochondrial RNA read content, data were then log-normalized, scaled and used for PCA dimensionality reduction on 2,000 variable features. Ten PCs were used for subsequent clustering and UMAP (Fig. S6a). Clustering with a 0.4 resolution identified 8 clusters: 0 and 2, composed of mainly SHH-induced samples, shared similar gene expression patterns and differed mainly in the n. of UMIs and housekeeping transcripts content (Fig. S6b), and were marked by neural progenitors markers, so they were annotated as SHH progenitors; 1 and 3 were similarly composed of control progenitors; 4 was marked by neuronal marker genes; 5 was mainly composed by low-UMIs cells with enriched ribosomal protein genes expression and was therefore annotated as low-quality and excluded; 6 was marked by extracellular matrix components and labelled as pericyte-like cells; 7 was composed by very few cells with extremely high UMI content and enriched nuclear/non-coding RNA markers, and removed for further analyses. After this second filtering, PCA and UMAP analyses were performed again on the subset to produce the plots shown in Fig. 4 and S6.

Gene set enrichment analysis on gene ontology terms and KEGG pathways was performed with the functions gseGO() and gseKEGG() in clusterProfiler v. 3.18.1 (Yu et al., 2012), with a minGSSsize set at 10, max set at 500, pvalueCutoff set at 0.05. Input to this analysis was a log fold-change-ranked list of differentially expressed genes computed with the FindMarkers() function in Seurat with min.pct and logfc.threshold both set at 0.1, between control and SHH-induced cells within the progenitors clusters 0-3.

Module scores shown in Fig. 4a and d were computed with the AddModuleScore function in Seurat, with the following genes: *SHH*, *FOXA2*, *NKX2-2*, *OLIG2*, *NKX6-1*, *NKX6-2*, *PTCH1*, *HHIP* in the SHH module, *FOXA2*, *NKX6-1*, *SHH*, *FERDL3L*, *ARX, LMX1B* in the FP, *NKX6-1*, *NKX2-2*, *NKX2-9* in the p3, *SP8*, *NKX6-1*, *OLIG2* in the pMN, *IRX3*, *IRX5*, *PAX6*, *DBX2*, *DBX1*, *SP8*, *NKX6-2*, *PRDM12*, *NKX6-1*, *FOXN4* in the p0-2, *MSX2*, *PAX3*, *OLIG3*, *IRX3*, *IRX5*, *PAX6*, *PAX7*, *GSX2*, *ASCL1*, *GSX1*, *GBX2*, *DBX2*, *DBX1*, *SP8* in the dp1-6, *LMX1A*, *MSX1*, *MSX2*, *PAX3*, *WNT1* in the RP modules (selected from Delile et al., 2019).

Spatial reconstruction shown in Fig. 4e-g was performed with novosparc (Nitzan et al., 2019, https://github.com/rajewsky-lab/novosparc), with alpha set to 0.5. Inputs were: a gene expression matrix composed of cells in clusters 0-3, additionally filtered for having > 1200 UMIs, a z-score scaled expression matrix of positional marker genes selected as follows, and a list of 1000 highly variable genes computed in Seurat from the clusters 0-3 and integrated with the 32 positional marker genes. Gene expression data and annotation metadata from a developing spinal cord atlas were downloaded from Delile et al. (2019), and filtered for cells annotated in one of the 13 dorsal ventral progenitor domains. The FindMarkers() function was used to compute marker genes for each domain, and an *a priori* set of known positional markers was complemented with the identified markers with highest fold-change, filtered for being expressed in the human data (the final list is shown in Fig. 4e). To control for the robustness of the reconstruction performed on all cells, we sampled 100 times 4,000-5,000 random cells and found highly similar results on the marker genes. To compare the novosparc reconstruction of human data with mouse developing spinal cord data, we generated a z-score scaled gene expression matrix for 1000 variable genes from the novosparc data and computed a correlation matrix with the z-score scaled mouse data, after filtering for genes present in both datasets, converted from human to mouse with an orthology table obtained from biomart and additionally filtered for unambiguous orthology.

### qRT-PCR primer pairs

For qRT-PCR measurements of target RNAs, we used the following forward and reverse primers.

ASCL1 fw: CTTCACCAACTGGTTCTGAGG

ASCL1 rv: CAACGCCACTGACAAGAAAGC

CDR1as fw: ACGTCTCCAGTGTGCTGA

CDR1as rv: CTTGACACAGGTGCCATC

STAT3 fw: AACATGGAAGAATCCAACAACGG

STAT3 rv: TCTCAAAGGTGATCAGGTGCAG

GAPDH fw: AAGGTGAAGGTCGGAGTCAAC

GAPDH rv: GGGGTCATTGATGGCAACAATA

HPRT fw: ACCCCACGAAGTGTTGGATA

HPRT rv: AAGCAGATGGCCACAGAACT

SHH fw: AAGGATGAAGAAAACACCGGAGCG

SHH rv: ATATGTGCCTTGGACTCGTAGTAC

BMP4 fw: GCTGCTGAGGTTAAAGAGGAAACGA

BMP4 rv: CACTCGGTCTTGAGTATCCTGAG

FOXA2 fw: CCGTTCTCCATCAACAACCT

FOXA2 rv: GGGGTAGTGCATCACCTGTT

FOXG1 fw: CACTGCCTCCTAGCTTGTCC

FOXG1 rv: TGAACTCGTAGATGCCGTTG

OLIG2 fw: CCAGAGCCCGATGACCTTTTT

OLIG2 rv: CACTGCCTCCTAGCTTGTCC

NKX2-2 fw: CCGGGCCGAGAAAGGTATG

NKX2-2 rv: GTTTGCCGTCCCTGACCAA

NKX6-2 fw: GAGGACGACGACGAATACAAC

NKX6-2 rv: GTTCGAGGGTTTGTGCTTCTT

### guide RNA sequences

For most luciferase knock-downs, we used the previously validated PS18 crRNA and non-targeting control (NT) (Abuddayyeh et al., 2016) for both Psp-Cas13b and CasRx, while the complementary sequence was cloned downstream of the Renilla luciferase reporter cassette in a psiCHECK-2 plasmid (Promega). The same target sequence was also cloned downstream of a TagRFP reporter cassette in an ePB-BSD-TT piggyback vector (see plasmids) for testing constitutive and light-inducible CasRx knock-downs. For the RSD tethering experiments, the 3’UTR was further swapped with another validated protospacer sequence (Cox et al., 2017), targeting the KRAS mRNA. The CDR1as crRNAs were designed on the CDR1as backsplice junction. The STAT3 mRNA crRNA sequence was taken from Konermann et al. (2018). All guide RNA sequences were cloned into the pr026 plasmid, carrying either the Psp-Cas13b or CasRx direct repeat with two adjacent BbsI restriction sites for guide cloning (see plasmids).

NT guide: GTAATGCCTGGCTTGTCGACGCATAGTCTG

PS18 guide (luciferase and TagRFP 3’UTR): CATGCCTGCAGGTCGAGTAGATTGCTGT

KRAS guide (luciferase 3’UTR): AAACTATAATGGTGAATATCTTCAAATGATTT

CDR1as PS1 guide: GTGCCATCGGAAACCCTGGATATTGCAGAC

CDR1as PS2 guide: CCATCGGAAACCCTGGATATTGCAGACAC

STAT3 guide: ATCACAATTGGCTCGGCCCCCATTCCCACA

For the light-inducible CRISPRa experiments, we used the Tet6 sgRNA spacer sequence reported below, targeting CaSP1/2 and GAL4/UAS promoters. A sgRNA plasmid without any spacer cloned was used as a non-targeting guide control. We report below also all the tested guide RNA sequences for SHH and BMP4, designed after the Calabrese library (Sanson et al., 2018). All guides were cloned into the psgRNA2.0 plasmid carrying the SpCas9 sgRNA scaffold with two MS2 aptamers (Nihongaki et al., 2017, see plasmids).

Non-targeting guide: GAACGACTAGTTAGGCGTGTA

ASCL1 guide: GCAGCCGCTCGCTGCAGCAG

Tet6 guide: GTCTTCGGAGGACAGTACTC

SHH guide 1: CATCAGAAGACAAGCTTGTG

SHH guide 2: AAAAAACGTAGTCTTCTTCA

SHH guide 3: TTTCCTAAGATAAAGGTGGG

BMP4 guide 1: CTCGCTCGCCTCCCTTTCTG

BMP4 guide 2: GGGGCTCCCATCCCCAGAAA

BMP4 guide 3: GCCTGCTAGGCGAGGTCGGG

