## Supplementary material for "Spatio-temporal, optogenetic control of gene expression in organoids"

Ivano Legnini et al.

#### List of supplementary items

Supplementary note (p. 2)

Supplementary Figures 1-7 and legends (p. 7)

Supplementary table 1 (separate file)

Proteomic (LFQ and iBAQ values for 3 biological replicates times 2 technical replicates per condition along with annotations and statistics; conditions are non-targeting guide RNA, *NT*, and RFP-targeting guide RNA, *RFP* or *PS18*; see methods for more details on the analysis) and transcriptomic data (DeSeq2 output from the PiGx pipeline for 2 biological replicates per condition; see methods for more details on the analysis) for HEK293T cells where RFP was knocked down with CasRx.

Supplementary table 2 (separate file)

List (and information on availability) of the plasmids used in this work.

Supplementary video 1 (separate file)

SHH induction in hiPSCs: a laser scanning setup was used to stimulate SHH expression with the PA-Cre/Lox system (De Santis et al., 2021) in a squared region of interest at the center of a larger field of view. On the left, the RFP and GFP channels for the entire FOV at the end of induction; on the right, the photostimulated ROI in the GFP channel. Stimulation was performed overnight (ca. 16 hours).

Supplementary video 2 (separate file)

SHH induction in neural organoids: induction performed as in video S1. Four embryoid bodies are arranged in a dish and embedded in matrigel in order to use a single ROI to stimulate a limited region of all of them at once. On the left, the RFP and GFP channels for the entire FOV at the end of induction; on the right, the photostimulated ROI in the GFP channel. Stimulation was performed overnight (ca. 18 hours).

**Supplementary note: light-inducible knock-down of reporter and endogenous transcripts.**

Introduction

While gene activation based on light-inducible transcription (Polstein et al., 2015, Nihongaki et al., 2015a, Nihongaki et al., 2017, De Santis et al., 2021) has been effectively used in cell culture and even in organotypic culture and in vivo (Yamada et al., 2018), spatio-temporal control of RNA knock-down poses several unmet challenges.

Previous efforts in this direction combined light-responsive protein modules (Kennedy et al., 2010, Renicke et al., 2013, Kawano et al., 2015) with CRISPR/Cas9 (Nihongaki et al., 2015b, Zhou et al., 2017), to induce irreversible genetic mutations. However, the results can be difficult to interpret due to mutational heterogeneity and limited efficacy. An alternative approach, based on light-inducible knock-downs, has only been reported in cell lines, for example with photo-caged oligonucleotides/siRNAs (e.g. Mikat et al., 2007), or more recently with a genetically encoded optogenetic RNA interference (RNAi) system (Pilsl et al., 2020). While these methods are efficient and allow for precise temporal control in cell lines, their efficacy is unknown in more complex tissues and may be limited. For example, the RNAi approach used in Pilsl et al., is constitutively active in the dark and is inactivated by photo-stimulation, making it challenging to spatially control the knock-down of a given target in an organoid or *in vivo*. Furthermore, RNAi in general is known to suffer from off-target effects.

We therefore explored two alternative strategies for achieving optogenetic activation of CRISPR/Cas13, a relatively new molecular tool for efficient and programmable RNA targeting, by either controlling its transcription with photo-stimulation, or its enzymatic activity.

RNA knock-down by light inducible transcription of CRISPR/CasRx

We first tested the efficacy of constitutively expressed or light-inducible CasRx in knocking down reporter and endogenous transcripts (Fig. S2a) and here report our results. Constitutively expressed CasRx was able to efficiently knock-down a Tet-ON RFP reporter with a single guide RNA (PS18 adopted from Abudayyeh et al., 2016; hereafter referred to as “RFP guide RNA”; Fig. S2b). However, GFP (which tags the CasRx cassette) was also depleted for reasons that we do not fully understand (Fig. S2c). This effect was not restricted to GFP only, but also CasRx was depleted when targeting RFP (Fig. S2d), suggesting that such cross-talk occurs at the RNA level, as CasRx and GFP are co-expressed within the same cistron. GFP depletion upon RFP targeting seems an effect restricted to these two genes, since targeting a luciferase mRNA carrying the same target sequence with the same guide RNA did not induce any GFP nor CasRx depletion and the guide RNA targeting RFP did not induce any GFP nor CasRx knock-down in absence of RFP (Fig. S2d). On the other hand, exchanging the GFP tag with a different fluorescent protein (UnaG) retained this effect (Fig. S2e). However, this off-targeting seemed limited to the overexpressed cassette. In fact, to examine the occurrence of global off-targeting effects, which have been recently reported in certain conditions (Ai et al., 2021, Shi et al., 2021), we performed genome-wide mass-spectrometry-based quantitative proteomics. The only proteins with a statistically significant fold change greater than 2 when comparing cells transfected with a non-targeting guide (NT) vs. an RFP-targeting guide were RFP, CasRx and GFP (Fig. S2f). We also sequenced total RNA and found only six differentially expressed transcripts: four were highly homologous to 18S ribosomal RNA (a typical artifact of ribodepletion) and the remaining two were RFP and CasRx-GFP, with only RFP having a fold-change larger than 2 (Fig. S2g). Therefore, genome-wide protein and RNA quantification demonstrated the specificity of our knock-down approach in the tested conditions and within the analyzed time frame.

Given the targeting efficacy and specificity of the constitutive CasRx, we tested whether a light-induced CasRx is also capable of specifically knocking down RFP. We transfected HEK cells with the SCPTS CasRx systems together with either a non-targeting or an RFP-targeting guide and observed RFP knock-down efficiencies of 40-60% in the induced (lit) state, with residual activity of 10-30% in the uninduced (dark) state, depending on the construct (Fig. S2h). As before, GFP was unexpectedly depleted when CasRx was programmed to target RFP (Fig. S2i). We then tested the CaSP2-CasRx system, which provided a good trade-off between targeting efficacy and leakage, on endogenous targets. We designed two guide RNAs complementary to the circular RNA CDR1as and adopted a previously published sequence for the STAT3 mRNA (Konerman et al., 2018) and validated them with a constitutive CasRx (Fig. S2j). We transfected one CDR1as guide and the STAT3 guide together with the SCPTS-CaSP2-CasRx system, stimulated the cells with blue light for 24-36 hours and performed qRT-PCR for target quantification. As shown in Fig. S2k, we achieved ~71% and ~45% knock-down for CDR1as and STAT3 respectively, with ~24% and ~11% leakage.

We also generated two stable lines carrying the light inducible Cre/Lox CasRx system, with or without a CDR1as targeting guide (PS2 from Fig. S2j), to test the knock-down properties of this system. Upon doxycycline and blue light stimulation, we observed on average ca. 50% knock-down of CDR1as, with ca. 25% leakage with doxycycline treatment and no leakage without (Fig. S3l).

In summary, we found that CasRx-mediated knock-downs could be induced upon photo-stimulation, with a tradeoff between activity and background. For example, the CRISPRa system combined with our CaSP1 promoter was very effective upon light stimulation, but it also had high background. CaSP2 had lower efficiency, but also lower leakage. This is common for inducible systems, and one has to adjust the experimental conditions to reach a predetermined goal. In this case, one can titrate the minimum amount of CasRx required for efficient knock-down of a given target, and then adjust the experimental setup to have the lit state exceeding that threshold. These systems enabled silencing of reporter and endogenous targets with varying efficiencies and leakage. Further improvements may be possible, and similar approaches have been recently described (e.g. Blomeier et al., 2021). On the other hand, recent studies suggested that certain Cas13-mediated knock-downs can be prone to nonspecific off-target effects (Ai et al., 2021, Shi et al., 2021), which we did not observe in our experimental conditions but should be kept in mind when designing experimental setups and readouts.

<u>Attempts at Cas13 protein engineering for optogenetic RNA targeting.</u>

We also attempted at engineering Cas13 proteins themselves for making their targeting activity light responsive, rather than their synthesis.

We first optimized a luciferase assay for assessing the targeting efficiency of Cas13 in human cells (Fig S3a). We designed several guides for PspCas13b (Cox et al., 2017) and obtained high knock-down efficiency for all of them (Fig. S3a). We also designed a catalytically inactive version of PspCas13b (dCas13b) fused with the RNA silencing domain (RSD) of known RNA decay factors such as GW182 (TNRC6A and C, involved in microRNA-mediated targeting) and the endonuclease SMG6, which have been previously used for tethering experiments (Chen et al., 2009, Nicholson et al., 2014). Of these constructs, only the GW182 fusions were able to induce 50-60% luciferase knock-down, and only when targeting the 3’UTR (Fig. S3b). We further confirmed this finding by switching the target

3'UTR with a different sequence and observed similar knock-down efficiency with the TNRC6A/C fusions (Fig. S3c).

Another Cas13 effector, CasRx, induced strong luciferase knock-down, while its catalytically inactive form did not (Fig. S3d).

To produce a light-inducible Cas13 system, we designed ten split PspCas13b pairs, which we fused with two distinct photo-dimer pairs (CRY/CIB and pMan/nMag) that should mediate the reconstitution of the full-length protein upon light stimulation (Kennedy et al., 2010, Kawano et al., 2015, Fig. S3e). For CasRx, we designed 14 split pairs and fused them with one pair of photo-dimers. We also split dCas13b as well as dCasRx and the GW182 RSDs and fused them in different arrangements with the photo-dimers (Fig. S3e).

We transfected all the split constructs together with the luciferase reporter and guide RNA and tested the targeting efficiency upon blue light stimulation for 24 hours (Fig. S3f). While light stimulation per se did not affect *wt* Cas13 knock-down efficiency, most of the Cas13b split pairs retained strong activity but reacted to light stimulation in an unexpected way, with the target being efficiently knocked down in the dark state, and the effect being reverted to different extents in the lit state (Fig. S3g-j). Although we do not have an exhaustive explanation for this observation, we hypothesize that the split protein is still able to correctly fold in the dark state, while the change of conformation induced by photo-dimerization constrains target recognition, enzymatic activity and/or protein stability in the lit state, as observed in a different context in Zetsche et al., 2015. For the pMag/nMag PspCas13b-763 pair, we validated the initial screening result with additional independent replicates and confirmed that the light switch can induce up to 10-fold knock-down of the luciferase reporter. We note however that the system is leaky, with approximately 50% knock-down happening in the lit (here to be considered the "inactive") state.

In addition to the photodimerization/phototethering approaches, we also devised a few single-chain designs and a few split designs combined with "inverted" photodimer modules such as pdDronpa1 and LOVTRAP, which should dissociate in response to blue or cyan light stimulation (Zhou et al., 2017 and Wang et al., 2016). We fused two pdDronpa1 modules to Cas13b N- and C-termini, in a way that dimerization would cause steric hindrance at the interface with the guide:target RNA. With this approach, we obtained strong luciferase knock-down upon cyan light stimulation, but it was also very leaky with more than 60% knock-down in the dark state (Fig. S3k). The same design for CasRx did not produce strong target knock-down (Fig. S3k). We tried to use the functional Cas13b-763 split pair in combination with the LOVTRAP photodimer pair, but we still had strong target knock-down in the dark state (Fig. S3k). Finally, we inserted a LOV2-Ja module in a supposedly flexible region of CasRx (Zhang et al., 2018) and in the Cas13b-763 split site, reasoning that the rigid structure of the protein module would impair RNA targeting in the dark state, and its relaxation upon blue light stimulation would allow the protein to fold and acquire an active state. Again, we observed a strong knock-down in the dark state for Cas13b, and no significant activity of CasRx (Fig. S3k).

All these photo-stimulation experiments were performed with a LED board which can accommodate a 96-well plate within a cell culture incubator. Each of the 96 LEDs can be programmed independently and illuminate one single well with the desired ON/OFF pattern over time. We report the details for reproducing such device in the methods section and in a public repository (https://github.com/BIMSBbioinfo/casled).

From these experiments, we conclude that: I) Cas13b can be efficiently split to produce two protein modules which will naturally assemble and reconstitute the full-length functional protein; II) Cas13b can be fused with RNA silencing domains from GW182/TNRC6 proteins and be programmed to repress target genes by binding their 3'UTR; III) light-inducible target repression of engineered Cas13 proteins cannot be achieved with most of our designs, and when achieved it follows different principles and kinetics than expected. We note that points I) and II) may be of interest for synthetic biology applications. We found several functional split sites for Psp-Cas13b, which could be used for example for conditional knock-downs where the two protein modules are expressed from different promoters (as in Kempton et al., 2020), or for fitting the Cas13 cassette into a compact viral vector (as in Chew et al., 2020). Also, we found that we can achieve target interference using a mutated Psp-Cas13b fused with domains of GW182 proteins, the effectors of microRNA silencing. Interestingly, the extent of silencing (ca. 50%) was consistent with that of microRNAs, and it worked only when tethering occurred on the 3'UTR, the natural platform for microRNA binding. These constructs might be used to manipulate gene expression or to study GW182 silencing activity.

### Figure S1

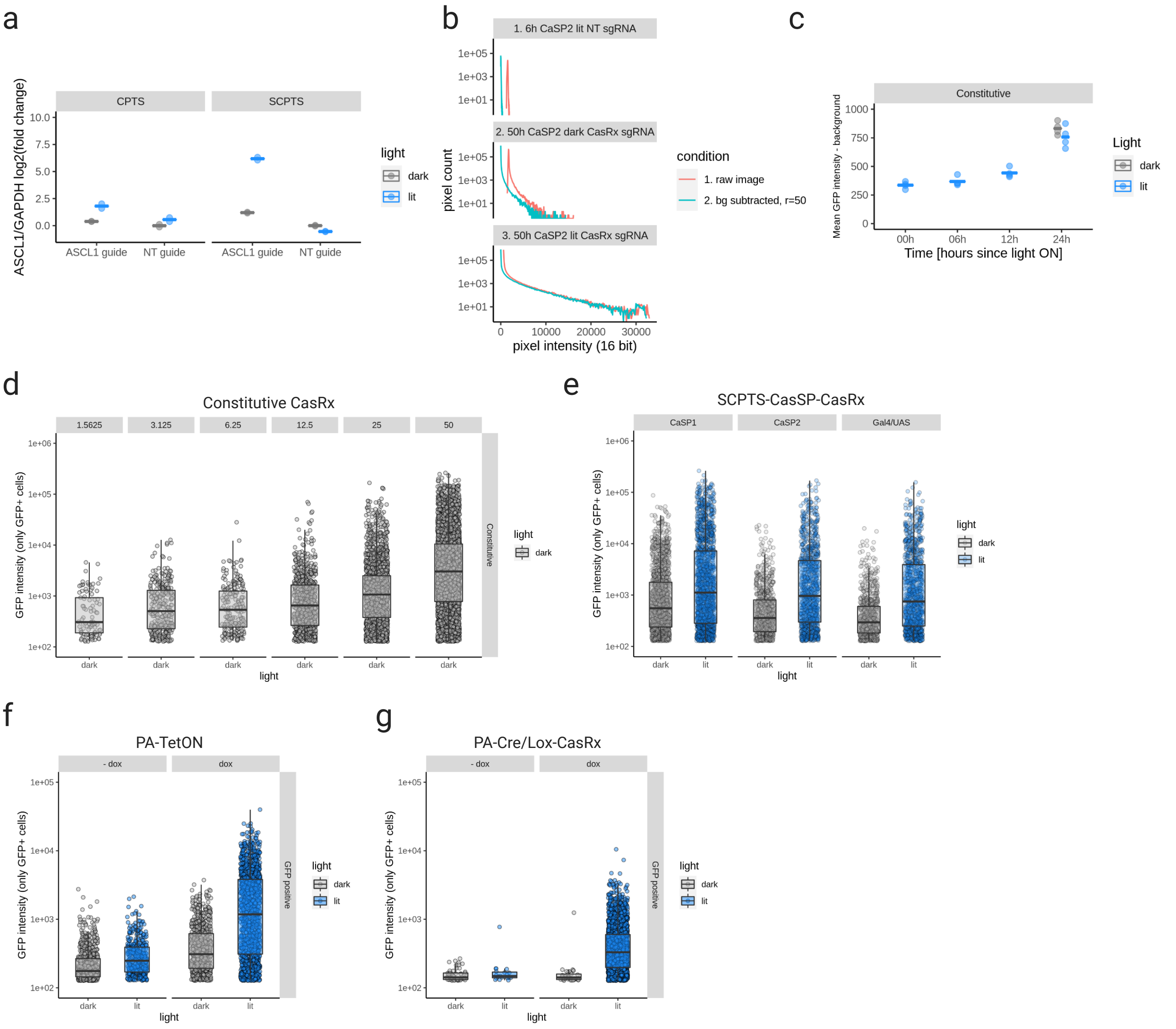


**Supplementary Figure 1. Light-inducible gene activation and knock-down modules (related to Figure 1 and the Supplementary Note).**

**a.** qRT-PCR measurement of ASCL1 mRNA induced with the CPTS (left) and SCPTS (right) systems (Nihongaki et al., 2017), combined with either a non-targeting sgRNA (NT) or an ASCL1 promoter-targeting sgRNA (ASCL1 guide), in dark or lit (24h pulsed illumination) conditions. mRNA level is plotted as Delta-delta Ct against GAPDH as reference gene and the NT dark condition as reference sample. Experiment was performed in two replicates (each dot). **b.** Histograms of pixel intensity with and without background correction (green and red), performed with the subtract background function in Fiji/imageJ with a rolling ball radius of 50, for three representative samples of the SCPTS CaSP2 CasRx system (top: 6h time point with non-targeting guide, middle: 50h time point with the Tet6 targeting guide in the dark, bottom: 50h time point with the Tet6 targeting guide with illumination). **c.** Background-subtracted mean GFP intensity is plotted against the selected time points, in either dark or lit conditions, for a constitutively expressed CasRx cassette (under a EF1a promoter). The experiment was performed in four replicates, each dot represents a replicate and horizontal lines represent means over each condition. **d-g.** FACS analysis of GFP in HEK293T cells transfected with a titration of a constitutive (d) CasRx plasmid (nanograms are indicated on the top of each panel), the SCPTS CaSP1/2 and Gal4/UAS CasRx system (e), and stable lines with the PA-TetON (f) and the Cre/Lox (g) system with or without doxycycline. Cells were gated into GFP-negative and positive to better compare GFP levels in transfected cells, which are shown here. 10000 total events were measured per condition, and boxplots with fluorescence quartiles are overlaid to each distribution (grey or blue for dark and lit conditions).

### Figure S2

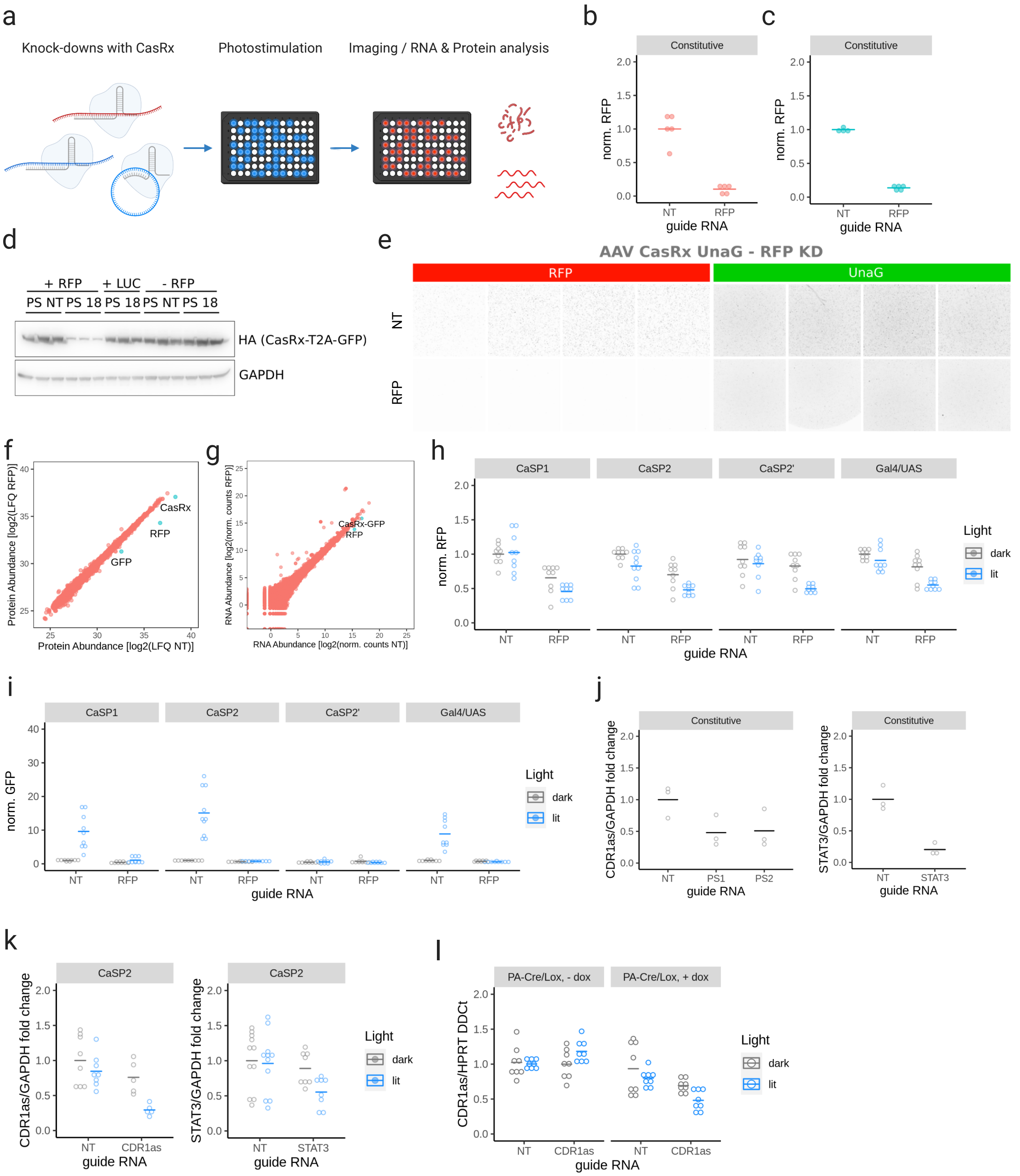


**Supplementary Figure 2 (related to figure 1 and the Supplementary Note). Light-inducible RNA knock downs with optogenetic activation of CasRx transcription.**

**a.** CasRx knock-downs of reporter and endogenous target RNAs, along with readouts of efficiency, off-targeting and leakage. **b-c.** Background-subtracted RFP and GFP signal intensity normalized on the non-targeting guide control (NT), with a constitutively expressed CasRx cotransfected with the RFP reporter, a non-targeting guide or an RFP-targeting guide (RFP). Horizontal bars: mean of all replicates (4-5). **d.** Western blot analysis of HEK cells transfected with a constitutively expressed CasRx, along with either a non-targeting

(NT) or RFP-targeting (PS18) guide, in presence or absence of the RFP reporter (+/- RFP) or in presence of a Luciferase reporter carrying the same binding site for the RFP-targeting guide (+ LUC). Western blot was stained for the HA tag present in the CasRx protein and for GAPDH as a loading control, as indicated on the side. The experiment was performed in triplicate. **e.** Fluorescence microscope images for RFP (left) and UnaG (right), in HEK cells cotransfected with a constitutively expressed CasRx tagged with a UnaG fluorescent protein, the RFP reporter, and either a non-targeting (NT) guide or an RFP-targeting guide (RFP). The experiment was performed in four replicates. **f.** Protein abundance (average over 6 replicates: three biological times two technical) in cells transfected with a constitutively expressed CasRx, the RFP reporter, a non-targeting guide (NT) or an RFP-targeting guide (RFP). Protein abundance was measured by shot-gun proteomics. LFQ: log2 Label-Free Quantification intensity. Green: proteins with statistically significant and > two-fold change. **g.** RNA abundance (average over two replicates) in cells transfected with a constitutively expressed CasRx, the RFP reporter, a non-targeting guide or an RFP-targeting guide (RFP). RNA abundance is expressed as normalized transcript counts as from the STAR+Deseq2 output in the PiGx analysis pipeline. Highlighted in green the only two transcripts with statistically significant difference between the two conditions, excluding four transcripts with high homology to the 18S rRNA interpreted as artifacts of the ribodepletion protocol. **h-i.** As in b and c, for the light-inducible promoters indicated on top of each panel used in combination with the SCPTS system. Grey and blue indicate dark and lit conditions, horizontal bars indicate mean of all replicates (9 for all, 10 for one condition) for each condition. For RFP, CaSP1, CaSP2 and CaSP2' (same as CaSP2 but with a CasRx devoid of the GFP tag) had a significant difference in fluorescence intensity between dark and lit conditions for the RFP-targeting guide (p-value $< 0.05$, Student's t test). All systems had non-significant difference between dark and lit conditions with the non-targeting guide. **j.** qRT-PCR data for CDR1as (left) and STAT3 (right), where GAPDH was used as a reference mRNA and the non-targeting guide control as the reference sample for normalization, for cells treated with a constitutively expressed CasRx along with either a non-targeting guide (NT) or one of two CDR1as-targeting guides (PS1 and 2), or a single STAT3-targeting guide (STAT3). p-value $< 0.05$ for NT vs PS1/2 CDR1as and for NT vs STAT3, Student's t test. **k.** qRT-PCR for CDR1as and STAT3, normalized to GAPDH and the non-targeting guide control as the reference sample for normalization, for cells treated with the CaSP2 CasRx light-inducible system along with either a non-targeting guide (NT) or one CDR1as-targeting guide (CDR1as), or a STAT3-targeting guide (STAT3). Grey, blue colors: dark, lit. Horizontal bars: mean over each condition. p-value $< 0.05$ in dark vs lit, Student's t test. **l.** As in j, in a stable cell line produced with the Cre/Lox CasRx system with either no guide or one CDR1as guide (PS2), in the dark and lit conditions and with or without doxycycline treatment. p-value $< 0.05$ for the CDR1as PS2 in dark vs lit conditions, Student's t test.

### Figure S3

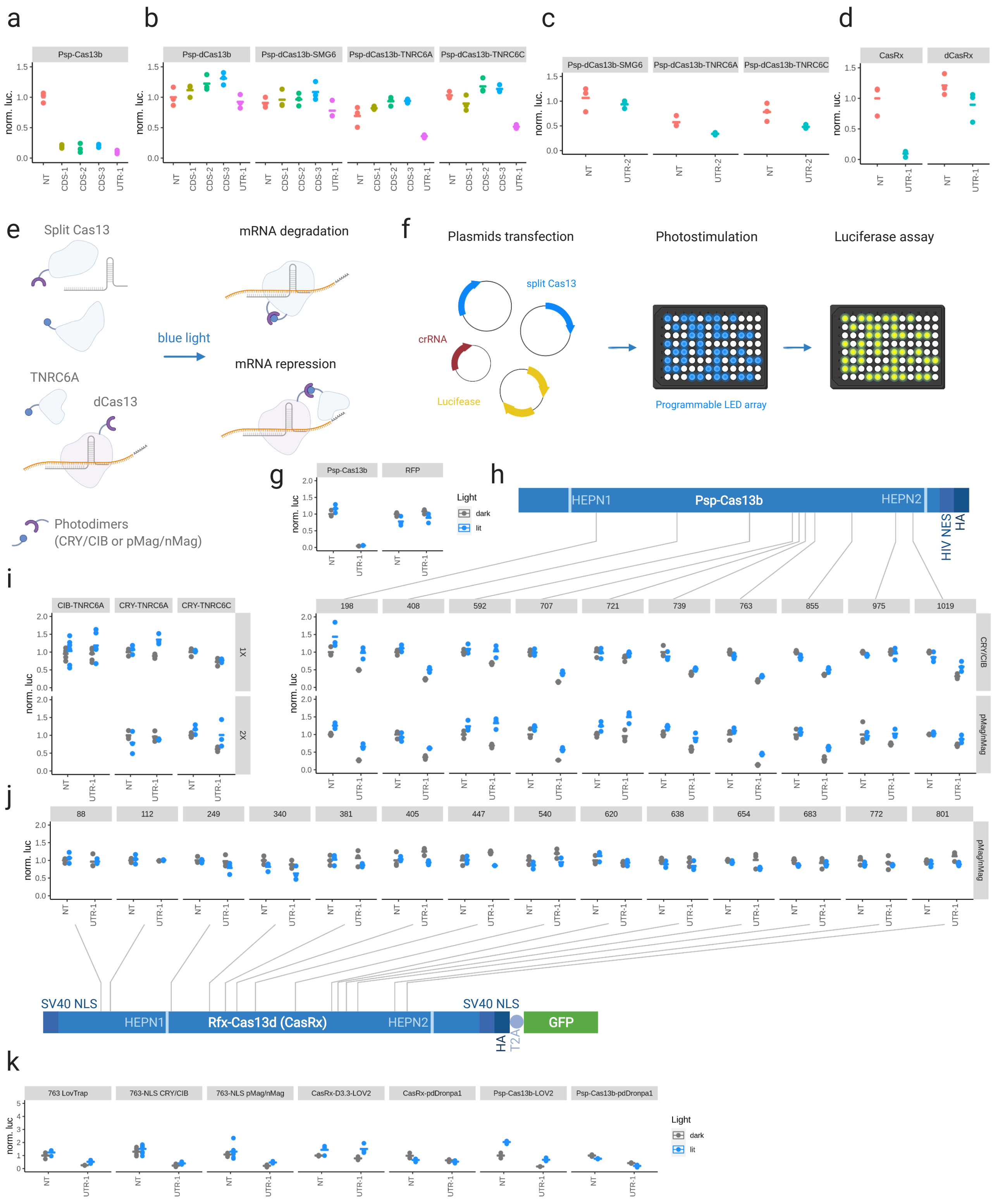


**Supplementary Figure 3 (related to the Supplementary Note). Attempts at engineering a light-inducible Cas13 protein.**

**a.** Normalized luciferase activity (Renilla/Firefly, non-targeting guide control as a reference sample) from HEK293T cells transfected with Psp-Cas13b, a luciferase reporter, and either a non-targeting guide (NT) or one of 4 Renilla luciferase-targeting guides, targeting the coding sequence (CDS1-3) or the 3'UTR (UTR-1). Horizontal bars indicate the mean for all replicates. **b.** As in a., but with one of four catalytically inactive Psp-Cas13b (Psp-dCas13b) versions: one as it is, the others fused with an RNA-silencing domain from SMG6, TNRC6A or TNRC6C. **c.** As in b., using a luciferase reporter with a different 3'UTR and a guide RNA targeting the said 3'UTR (UTR-2). **d.** As in a., with CasRx and catalytically inactive CasRx (dCasRx), with either the non-targeting (NT) or the luciferase UTR-1 guide. **e.** Light-inducible Cas13 designs. Top: a split Cas13 used in combination with a pair of photodimers, to achieve light-inducible reconstitution of full-length Cas13 upon light stimulation. Bottom: a dCas13 and an RNA-silencing domain (RSD) used in combination with a pair of photodimers to induce RSD tethering on the target upon light stimulation. **f.** Experimental setup for testing all the light-inducible Cas13 designs: HEK293T are transfected in a 96-well plate format, which is placed on a LED array for 24h pulsed-light stimulation, and then the same plate is directly used for luciferase activity measurement. **g.** Positive (a *wt* Psp-Cas13b-expressing plasmid) and negative (an RFP-expressing plasmid) controls of the light-inducible Cas13 screening experiment, co-transfected with either a non-targeting (NT) or luciferase-targeting (UTR-1) guide. **h.** Schematic representation of the Psp-Cas13b protein sequence, with its 2 catalytic HEPN domains, a nuclear export signal (HIV NES), and C-terminal HA tag. The position of each split site is indicated as a vertical gray bar, which points to the corresponding luciferase data. Luciferase knock-down results are shown below for the Split Psp-Cas13b designs, where the two split parts of Psp-Cas13b were fused with the CRY2/CIBN (top) or pMag/nMag (bottom) photodimers. For each construct, the non-targeting dark control is set to 1 as a reference. Grey and blue colors indicate dark and lit conditions. **i.** Luciferase knock-down results for the phototethering designs with either 1 (top) or 2 (bottom) CIBN domains fused to Psp-dCas13b, and a CRY2 domain fused to the RNA-silencing domain from TNRC6a or C. **j.** Same as i., were CasRx was split and used in combination with pMag/nMag photodimers. **k.** Luciferase knock-down results for additional light-inducible constructs described in the Supplementary note. All experiments were performed in triplicate.

### Figure S4

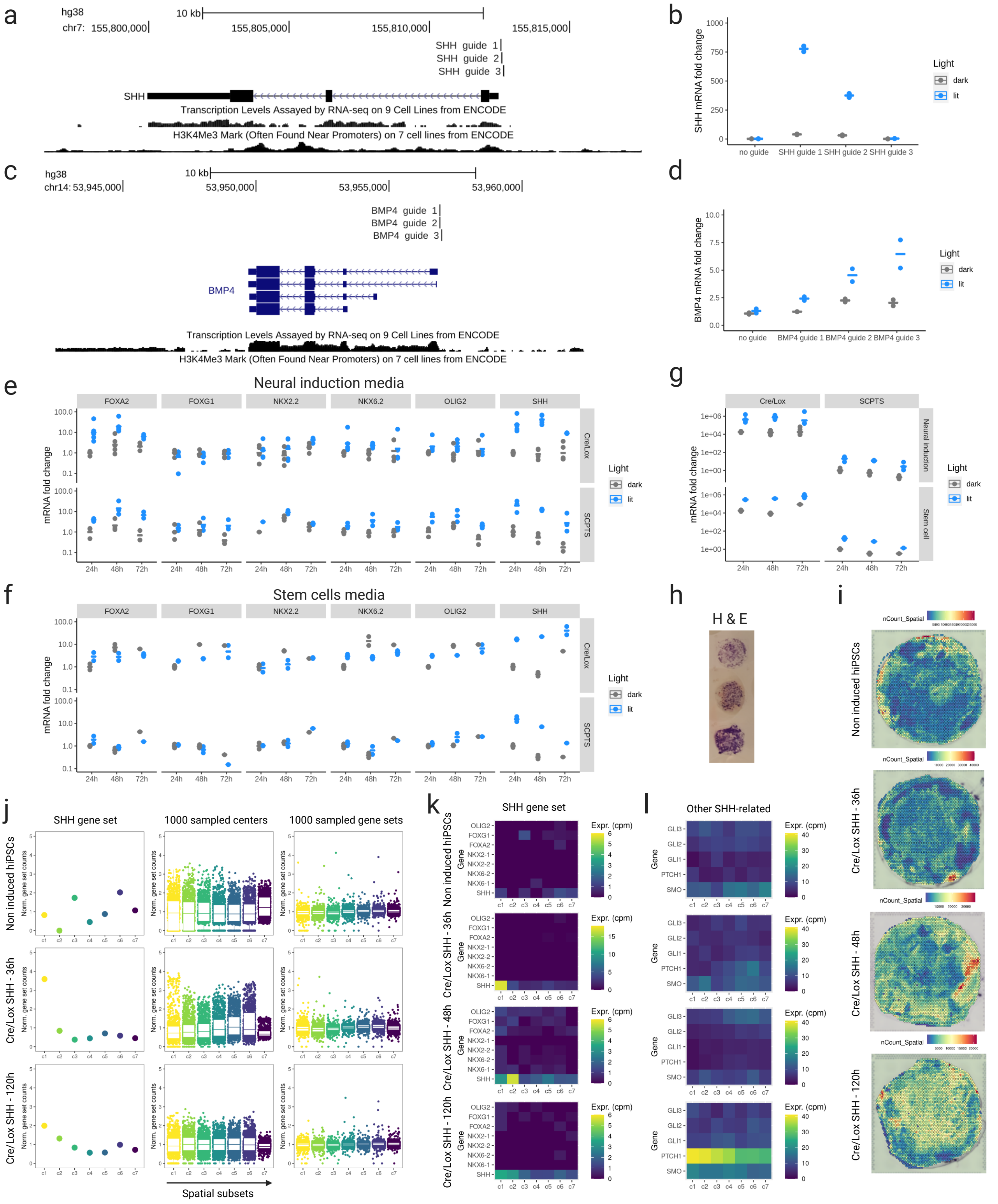


**Supplementary Figure 4 (related to figure 3). Optogenetic stimulation of Sonic Hedgehog in human stem cells.**

**a.** UCSC genome browser screenshot showing the human *SHH* locus along with transcription and H3K4Me3 mark for open chromatin available from ENCODE. The three guide RNA sequences we used for programming the light-inducible SCPTS dCas9 are shown in the top. **b.** qRT-PCR measurement of the *SHH* mRNA in

HEK293T cells transfected with the SCPTS system without guide or with one of the three *SHH* promoter targeting guides, with 24h light stimulation or in the dark. Values are fold change over the no-guide dark control sample, GAPDH was used as a reference gene. The experiment was performed in duplicate. **c-d.** As a-b, for *BMP4*. **e.** qRT-PCR measurement of the mRNAs indicated on top in hiPSCs induced for *SHH* expression with the SCPTS loaded with the *SHH* guide 1 or the Cre/Lox system (as indicated on the right), with or without light stimulation, for 24-72h. Data represent fold change over the 24h dark control. For SCPTS, *SHH*, *FOXA2*, *NKX6-2* and *OLIG2* had a p-value < 0.05 for the light term, calculated with the anova() R function on a linear mixed effects model with time and light as variables. Experiment was performed in five replicates; measurements were performed in 2-5 of the replicates depending on the sample. **f.** As in e, but with stem cell media instead of neural induction media. **g.** qRT-PCR measurement of the *SHH* mRNA using the SCPTS system loaded with the *SHH* guide 1 (right) or the Cre/Lox system (left), with or without light stimulation, for 24-72h. Data are the same as in e-f, but plotted as fold-change over the SCPTS dark 24h control for both series (SCPTS and Cre/Lox), to show the basal activation of the Cre/Lox system in the dark state. Experiment was performed in 5 replicates; measurements were performed in 2-5 replicates depending on the sample. **h.** Representative H&E staining on three replicates of hiPSCs cultured on a PET membrane and transferred onto a glass slide. **i.** UMI counts (color map) for the Visium capture areas in the 4 hiPSC Cre/Lox SHH samples at 0,36,48 and 120h of induction. **j.** Left: normalized counts of a *SHH* gene set (*SHH*, *FOXA2*, *FOXG1*, *NKX2-1*, *NKX2-2*, *NKX6-2*, *NKX6*-1, *OLIG2*), obtained by adding the counts of each gene in the set in each of the 7 concentric circles (c1-7 with c1 being the central one), normalized by the total transcript counts within each circle. Middle left: same as in the left, but sampling 1000 times a random central spot for drawing the concentric circles. Middle right: same as before, but sampling 1000 times a random gene set on the same concentric circles of the left panel. Right: heatmap showing transcript counts per million (cpm) for all genes in the *SHH* gene set across the 7 concentric circles (c1-7). Data shown here for hiPSCs Cre/Lox *SHH* induced for 0, 36 and 120h, while 48h is shown in Fig. 4e. The circles which resulted significant over sampling both centers and gene set were c1 at 36h, 48h and 120h and c2 at 48h (p-value < 0.05). **k.** Heatmap showing transcript counts per million (cpm) for the individual genes in the the *SHH* pathway across the 7 concentric circles (c1-7) and the 4 samples (induction of 0h, 36h, 48h and 120h). **l.** Same as k, for additional genes involved in the SHH pathway.

#### Figure S5

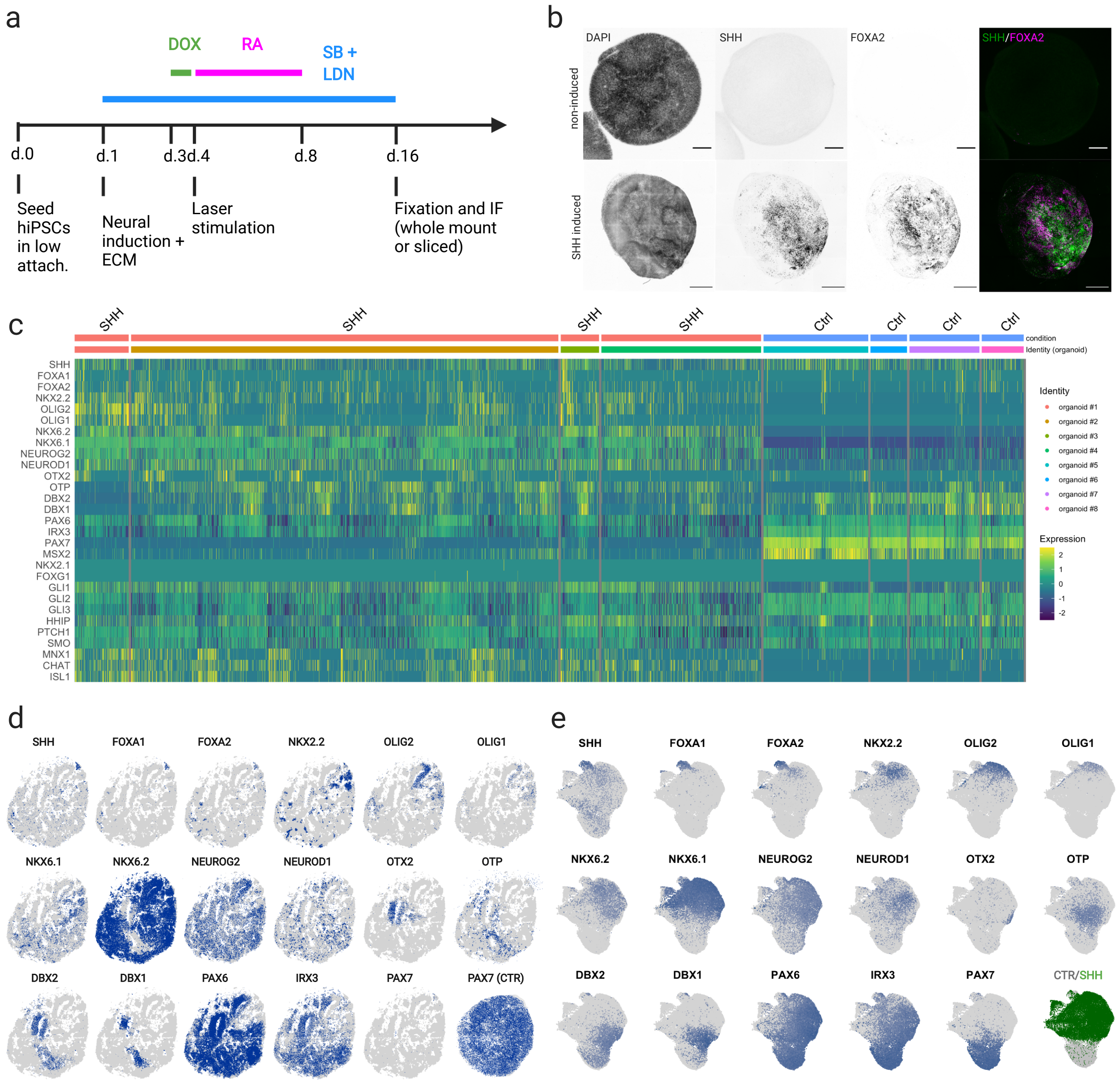


**Supplementary Figure 5 (related to figure 3). Optogenetic stimulation of Sonic Hedgehog in human organoids.**

**a.** Sketch of the protocol used for neural organoid derivation. **b.** Imaging of DAPI, *SHH* (NeonGreen fluorescent tag, labeled as GFP for simplicity) and *FOXA2* (immunofluorescence) in whole-mount neural organoids without induction (top) and with laser induction of *SHH* in the south-east pole. Signal is shown separately in grey scale for each target and merge for *SHH* and *FOXA2* in green and magenta (right). Scale bar: 100 $\mu$m and 500 $\mu$m for non-induced and induced organoids respectively. **c.** Heatmap of log-normalized transcript expression for a panel of transcripts of interest involved in the *SHH* pathway, quantified by Molecular Cartography spatial transcriptomics, for 8 organoids (4 controls and 4 SHH-induced as indicated on top). **d.** Molecular Cartography signal of selected transcripts of interest overlaid on a grey mask of a representative SHH-induced organoid cryosection. Bottom right: *PAX7* signal for a representative control organoid section. **e.** UMAP plot of log-normalized expression of the same transcripts as in e. for all detected cells in the 8 organoids. Lower right: organoids condition (control and SHH-induced in grey and green).

### Figure S6

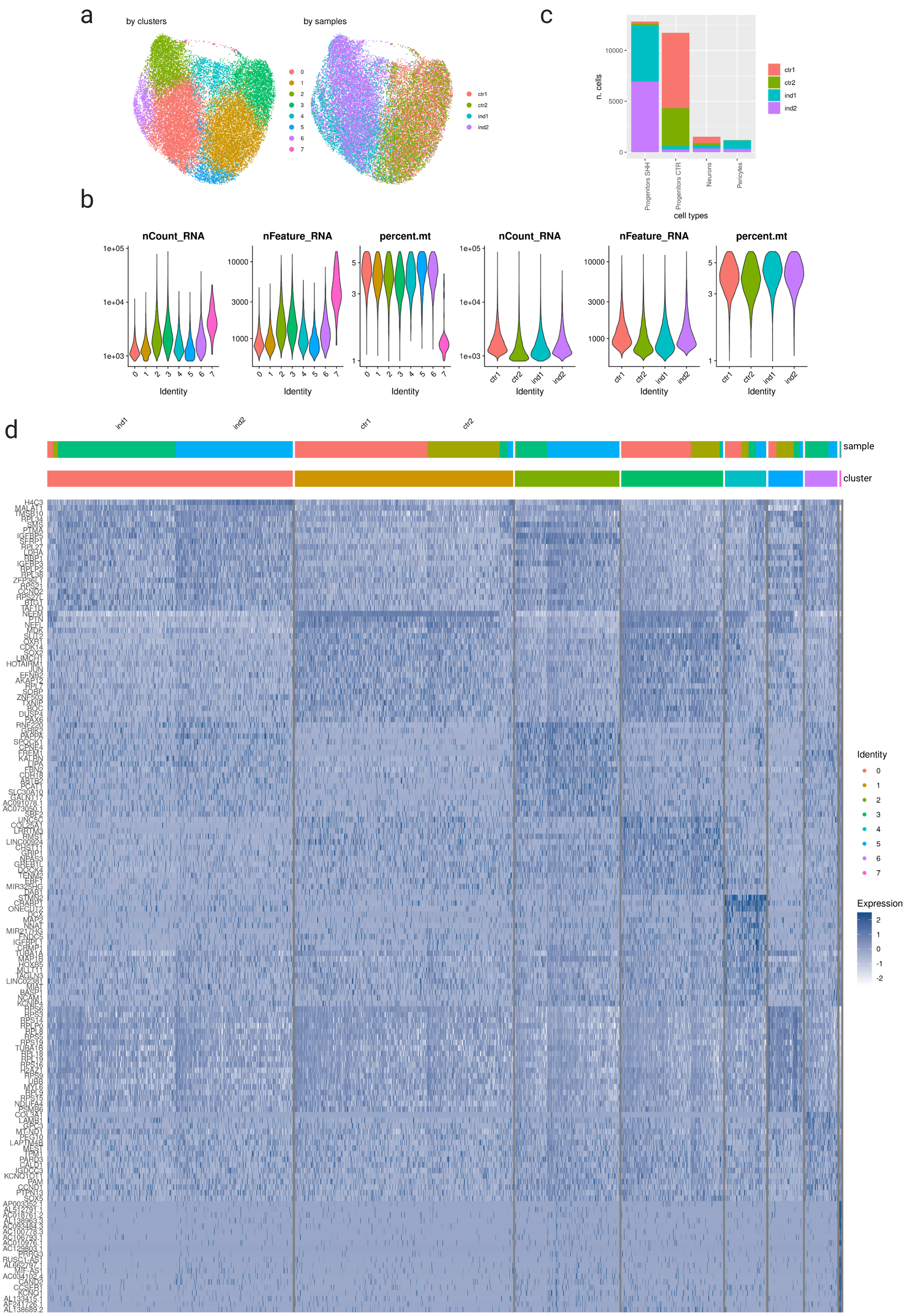
a
by clusters
by samples
0
1
2
3
4
5
6
7
ctr1
ctr2
ind1
ind2
c
n. cells
10000
5000
0
Progenitors SHH
Progenitors CTR
Neurons
Pericytes
cell types
b
nCount_RNA
nFeature_RNA
percent.mt
Identity
d
sample
cluster
Identity
Expression

**Supplementary Figure 6 (related to Figure 4). Molecular effects of SHH and spatial gene expression patterns in neural organoids.**

**a.** Identified clusters on all sequenced cells with > 800 UMIs and < 5% mtRNA, in UMAP space. **b.** and right: distribution of gene count (*nFeature*), UMI count (*nCount*) and mtRNA (*percent.mt*) by clusters and samples. **c.** Cell number by cell type and sample. **d.** Heatmap of top 20 marker genes expression for all clusters.

### Figure S7

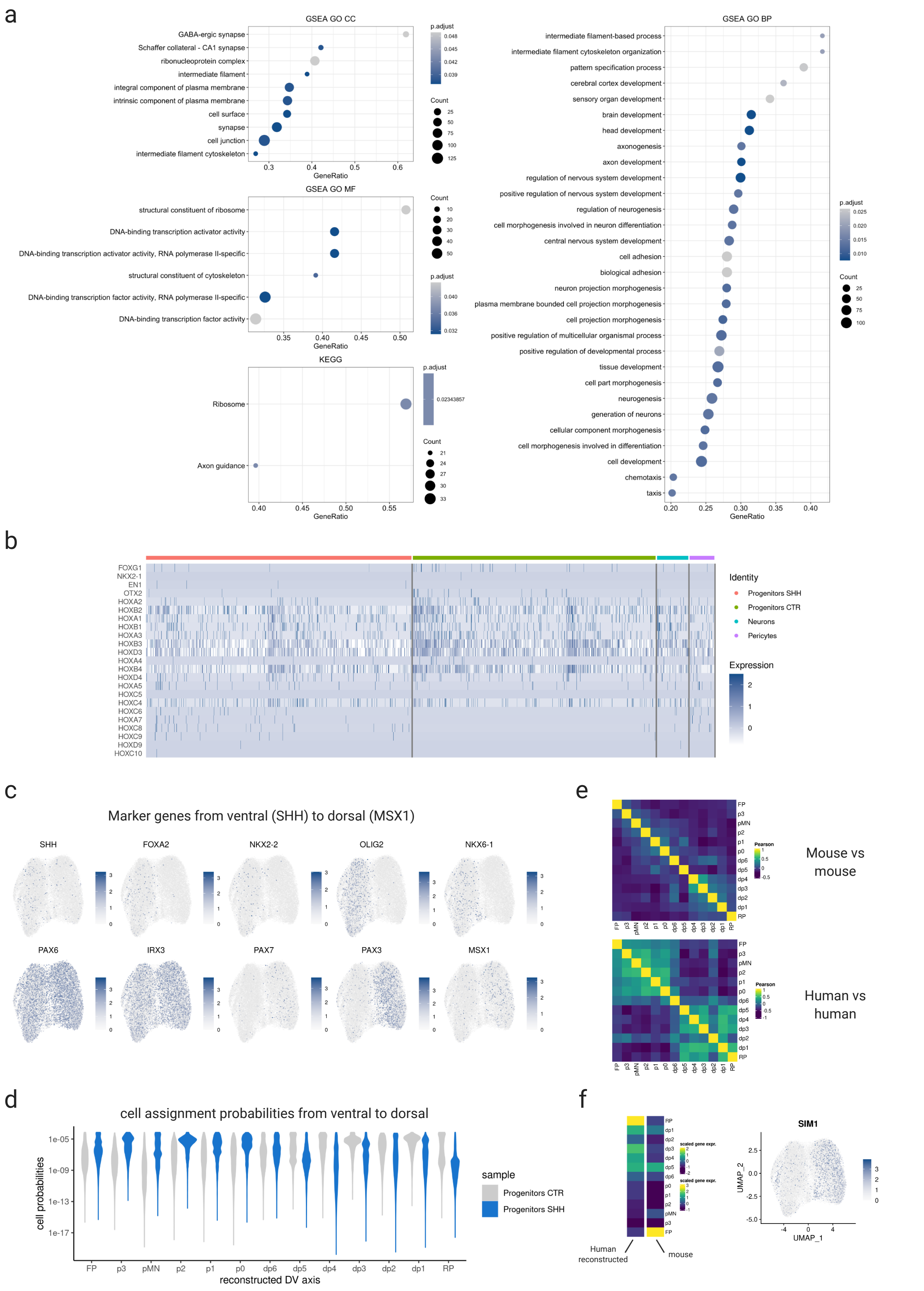
a
GSEA GO CC
GABA-ergic synapse
Schaffer collateral - CA1 synapse
ribonucleoprotein complex
intermediate filament
integral component of plasma membrane
intrinsic component of plasma membrane
cell surface
synapse
cell junction
intermediate filament cytoskeleton
GeneRatio
p.adjust
Count
GSEA GO MF
structural constituent of ribosome
DNA-binding transcription activator activity
DNA-binding transcription activator activity, RNA polymerase II-specific
structural constituent of cytoskeleton
DNA-binding transcription factor activity, RNA polymerase II-specific
DNA-binding transcription factor activity
KEGG
Ribosome
Axon guidance
GSEA GO BP
intermediate filament-based process
intermediate filament cytoskeleton organization
pattern specification process
cerebral cortex development
sensory organ development
brain development
head development
axonogenesis
axon development
regulation of nervous system development
positive regulation of nervous system development
regulation of neurogenesis
cell morphogenesis involved in neuron differentiation
central nervous system development
cell adhesion
biological adhesion
neuron projection morphogenesis
plasma membrane bounded cell projection morphogenesis
cell projection morphogenesis
positive regulation of multicellular organismal process
positive regulation of developmental process
tissue development
cell part morphogenesis
neurogenesis
generation of neurons
cellular component morphogenesis
cell morphogenesis involved in differentiation
cell development
chemotaxis
taxis
b
FOXG1
NKX2-1
EN1
OTX2
HOXA2
HOXB2
HOXA1
HOXB1
HOXA3
HOXB3
HOXD3
HOXA4
HOXB4
HOXD4
HOXA5
HOXC5
HOXC4
HOXC6
HOXA7
HOXC8
HOXC9
HOXD9
HOXC10
Identity
Progenitors SHH
Progenitors CTR
Neurons
Pericytes
Expression
c
Marker genes from ventral (SHH) to dorsal (MSX1)
SHH
FOXA2
NKX2-2
OLIG2
NKX6-1
PAX6
IRX3
PAX7
PAX3
MSX1
d
cell assignment probabilities from ventral to dorsal
cell probabilities
reconstructed DV axis
FP
p3
pMN
p2
p1
p0
dp6
dp5
dp4
dp3
dp2
dp1
RP
sample
Progenitors CTR
Progenitors SHH
e
Pearson
Mouse vs mouse
Human vs human
f
scaled gene expr.
Human reconstructed
mouse
SIM1
UMAP_1
UMAP_2

**Supplementary Figure 7 (related to Figure 4). Molecular effects of SHH and spatial gene expression patterns in neural organoids.**

**a.** GSEA analysis on GO CC, MF and BP terms, and on KEGG pathways, performed on a rank list of differentially expressed genes between control and induced progenitors. **b.** Heatmap of HOX genes and additional anterior-posterior markers showing that control and induced organoids have hindbrain/spinal cord identity, marked by expression peak between *HOXB2* and *HOXC4*. **c.** Featureplot of log-normalized expression of a set of positional marker genes along the ventral (*SHH*)-dorsal(*MSX1*) axis. **d.** Assignment probabilities of progenitor cells from control and induced organoids to each of the 13 progenitor domains specified by positional markers of a mouse spinal cord atlas. **e.** Correlation matrices for mouse and human reconstructed DV gene expression domains, computed on 1000 highly variable genes filtered for expression in human and mouse datasets. **f.** Left: DV scaled (z-score) expression of *SIM1* from human reconstructed and mouse DV domains. Right: featureplot of log-normalized expression of *SIM1*.
